# Early Brain Functional Connectivity Changes Induced by Antidepressants and Placebo

**DOI:** 10.1101/2025.08.22.671857

**Authors:** Xiaoyu Tong, Gregory A. Fonzo, Nancy B. Carlisle, Hua Xie, Yevgeny Berdichevsky, Corey J. Keller, Desmond J. Oathes, Charles B. Nemeroff, Yu Zhang

## Abstract

Major depressive disorder (MDD) is a common and debilitating condition with limited treatment precision. While brain imaging has linked neural features to MDD diagnosis and treatment response, the underlying circuits and their early modulation by treatment remain unclear. To examine changes in functional connectivity within the first two weeks of treatment, we analyzed two independent cohorts of MDD patients receiving antidepressants or placebo. Across patients, a visual-precuneus-thalamus network showed increased functional connectivity regardless of treatment arm or clinical outcome. Placebo response involved attention and striatal systems, while drug-specific effects were localized to the amygdala, mid-cingulate, orbitofrontal cortex, and cerebellum, emerging only in a subset of medicated patients. Notably, the responses of those without drug-specific changes can be predicted with a placebo response prediction model. These early functional connectivity changes reveal common and distinct mechanisms of treatment effects, offering insights that could inform more personalized interventions for MDD.

## Introduction

Major depressive disorder (MDD) is a major global health burden, affecting millions worldwide with resultant impaired quality of life. Despite its prevalence, MDD remains a highly heterogeneous condition with modest treatment efficacy. This is largely due to limited understanding of its underlying pathophysiology as well as the mechanisms of action of extant treatments. This necessitates the continued reliance on trial-and-error approaches in clinical care. Encouragingly, recent advances in machine learning applications in neuroimaging have provided unprecedented opportunities to identify brain-wide neural biomarkers for MDD^8, 9^. These efforts have yielded pre-treatment brain signatures for MDD diagnosis^1, 2^ and antidepressant response^3-5^, offering critical insights into the neurobiology of the disorder and its treatments. However, these studies also reveal that not all brain regions implicated in MDD pathology are targeted by antidepressants, and vice versa. Indeed, antidepressants may not alleviate depression symptoms by simply reversing the neurophysiological abnormalities in patients^10^. This discrepancy implies the involvement of intermediary neural circuits that translate treatment-induced brain changes into symptom relief. Consequently, the indirect action of antidepressants on symptom-relevant brain regions may limit the predictive power and clinical utility of current pre-treatment biomarkers, highlighting a critical need for research that directly examines the brain changes induced by antidepressant treatment to bridge this knowledge gap.

Previous studies have investigated such treatment-induced brain changes by measuring glucose metabolism^6^ and indices of various neurotransmitter-defined neural circuits^7^. For instance, placebo was shown to induce widespread cortical metabolic changes over six weeks^6^ while enhancing endogenous opioid release within one week^7^. Antidepressants induce metabolic changes in subcortical and limbic structures in addition to the placebo effect^6^, and their effects on serotonin and norepinephrine availability have been the central hypothesis of antidepressant pharmacology for decades^11^. However, these local profiles of brain changes do not fully capture the complex interactions across distributed brain systems. In contrast, functional connectivity (FC) offers a network-level perspective that may better capture the systemic impact of antidepressants. For example, one recent study revealed decreased FC within a thalamo-cortico-periaqueductal network after 10-12 weeks of serotonin-norepinephrine reuptake inhibitor (SNRI) treatment compared to placebo^12^. Nevertheless, it remains unclear if such early FC changes are induced by antidepressants, placebo, or factors unrelated to treatment such as sample variability, limiting insights into the pharmacological pathways of antidepressants and opportunities for adaptive treatment based on early response.

Functional magnetic resonance imaging (fMRI) is well-suited for detecting brain-wide FC changes^13^ due to its high spatial resolution and capacity to assess subcortical regions. Resting-state fMRI, in particular, captures spontaneous neural activity reflective of intrinsic brain organization, making it ideal for studying general treatment-induced changes independent of task-specific engagement^14^. However, fMRI data are often challenged by low signal-to-noise ratio (SNR)^15^ that may lead to low test-retest reliability in FC^16^, limiting the replicability and clinical utility of its findings^17^. This unfavorable observation complicates the robust detection of subtle treatment-induced changes amid measurement noise and systematic variance that may be meaningful but obscure treatment effects, necessitating analytic approaches that can effectively disentangle treatment effects from noise and other sources of variation.

To address these challenges, we sought to systematically characterize early FC changes (one or two weeks following treatment onset) induced by antidepressants and placebo and delineate their effects while minimizing sample variability, using two large, independent fMRI datasets comprising 386 MDD patients treated with selective serotonin reuptake inhibitors (SSRIs; sertraline or escitalopram) or placebo. We develop a novel machine learning framework that leverages multiple scan sessions at each time point to isolate genuine treatment-induced FC changes from fluctuations due to other sources (Fig. 1a,b). Through this approach, we first characterize the universal placebo effect, i.e. the changes shared across treatment arms and different levels of treatment responsiveness. We then identify early placebo-related FC changes predictive of symptom improvement. Next, we apply a contrastive learning strategy to extract sertraline-specific FC changes, distinguishing real drug effects from placebo effects (Fig. 1c). Remarkably, we find that only a subset of sertraline-medicated patients exhibits the drug-specific early FC changes, and we show that the sertraline-medicated patients without such drug-specific FC changes respond as if treated with placebo. Finally, using the escitalopram-medicated independent cohort, we demonstrate that both early placebo and drug-specific FC changes are generalizable across medications and datasets. Our findings fill a critical gap in understanding the neural dynamics of early antidepressant action in MDD, decompose treatment effects to real drug effects and placebo influences, and offer a new framework for stratifying treatment response. These results lay the groundwork for future studies into the neural circuits underlying antidepressant pharmacology and support the development of personalized and biologically informed approaches to MDD care.

**Fig. 1.**
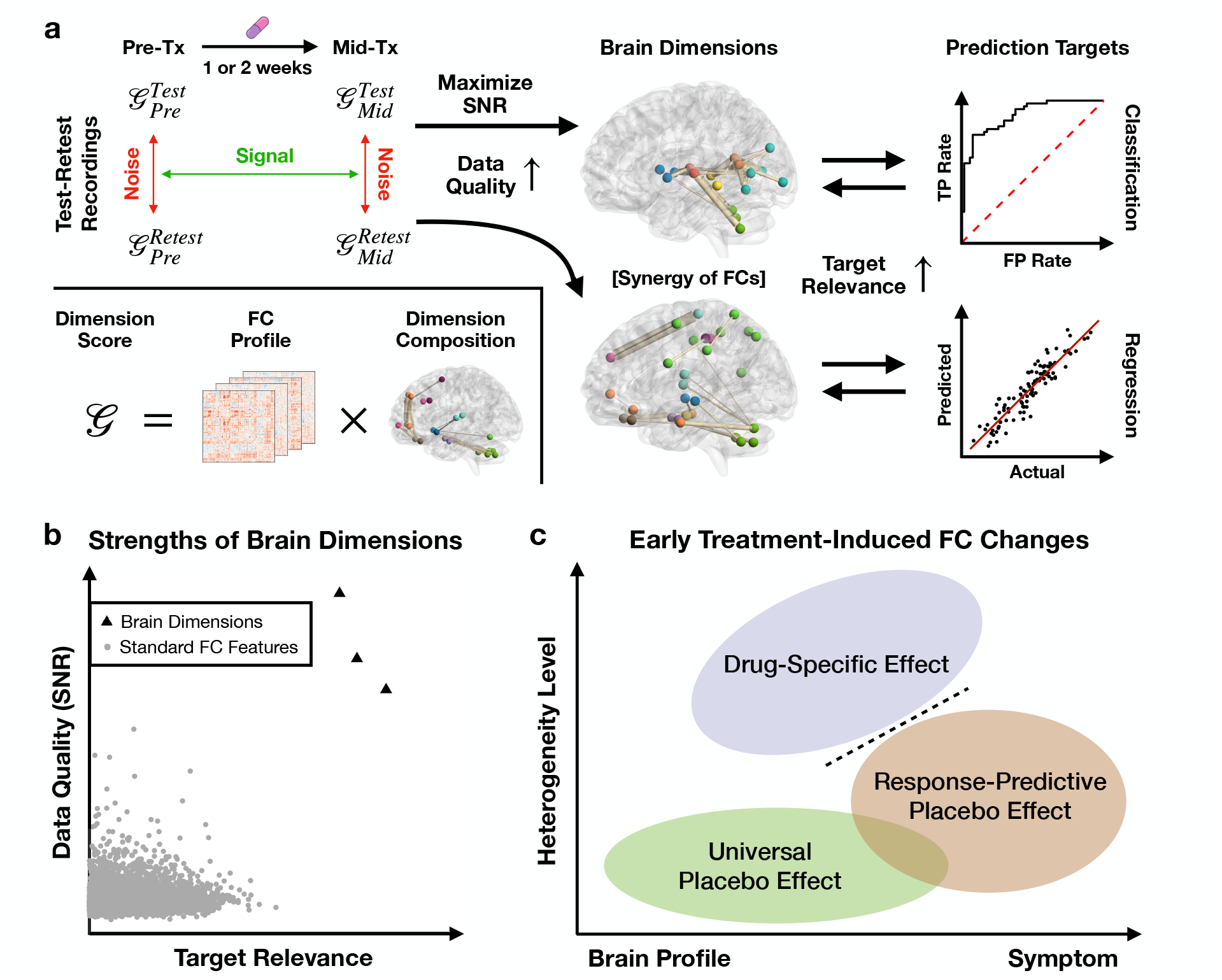
Identification of early treatment-induced FC changes. **a** Framework flowchart. Brain dimensions — weighted combinations of FC features — are constructed to enhance data quality, capture interactions across features, and increase relevance to specific prediction targets. Data quality is quantified by signal-to-noise ratio (SNR), where signal is defined as the mean difference in dimension scores between pre- and mid-treatment, and noise as the within-subject test–retest variability at the same time point. Each brain dimension score is a weighted sum of FC features, enabling synergy across features and improved interpretability. In the brain map, line width indicates the weight, and color indicates the network origin of each brain region. Dimensions are tailored to targets of interest, including categorical and continuous variables. **b** Strengths of brain dimensions. Compared to individual FC features, brain dimensions offer improved data quality and stronger relevance to prediction targets. **c** Facets of early treatment-induced FC changes. The *universal placebo effect* captures FC changes that are consistent across all medicated patients, regardless of drug type or clinical response, which may reflect changes in brain profile such as interregional communication efficiency and network topology. Yet, its magnitude may encode some heterogeneity of treatment response. The *response-predictive placebo effect* identifies placebo-driven FC changes that translate to symptom improvement, presumably mediated by treatment expectancy effects. The *drug-specific effect* denotes FC changes observed only in patients treated with active antidepressants, with no overlap with placebo-induced effects. Identified in an unsupervised manner, the drug-specific dimension reflects both brain profile and symptom-related changes presumably mediated purely by the mechanism of antidepressant action, revealing heterogeneity among patients that extends beyond differences in treatment response.

## Results

### Universal Placebo Effect

We first investigated whether antidepressant treatment induces shared changes in functional connectome across medicated MDD patients. Using a chronology prediction model trained to distinguish pre-treatment and post-initiation FC data in a within-subject pair (SFig. 1, **Methods**), we identified a brain dimension showing consistent FC changes after one week of treatment in both placebo and sertraline arms (placebo: accuracy = 94.4%, AUC = 0.991; sertraline: accuracy = 91.1%, AUC = 0.984, Fig. 2a,b). Notably, the brain dimensions derived from the placebo and sertraline arms were highly similar (cosine similarity = 0.813, SFig. 2a). Cross-arm testing confirmed the generalizability of these FC changes across treatment types (placebo-trained sertraline-testing: accuracy = 93.5%, AUC = 0.981; sertraline-trained placebo-testing: accuracy = 94.4%, AUC = 0.987). These findings suggest that the observed early FC changes may reflect a universal placebo effect and/or changes pertaining to repeated scan sessions. The use of each treatment arm as held-out test data for the other also demonstrated the generalizability of this brain dimension, with model performance further validated with permutation tests (p < 0.001 for both arms). Importantly, models trained on MDD patients failed to predict the chronology of untreated healthy subjects based on their one-week FC changes (placebo-trained: accuracy = 48.3%, p_binomial_ = 1; sertraline-trained: accuracy = 41.4%, p_binomial_ = 0.458), indicating that the changes observed in patients are less likely to reflect a main effect of repeated scanning. To ensure test-retest reliability, the models were trained on patients with two fMRI scans at both timepoints. We then tested whether these dimensions could generalize to 11 patients who had only one scan for at least one timepoint (6 in the placebo arm, 5 in the sertraline arm), thus also testing whether the model accurately predicts universal placebo-related changes independent of repeated scanning. Despite lower data quality, the two-scan-derived models showed outstanding generalizability to these patients (placebo- and sertraline-trained: accuracy = 100%, p < 0.0005), supporting the robustness and replicability of the identified FC change dimension as well as a high likelihood it is primarily detecting changes related to starting a new treatment irrespective of the type of treatment, namely a universal placebo effect, and not changes pertaining to repeated scanning.

**Fig. 2.**
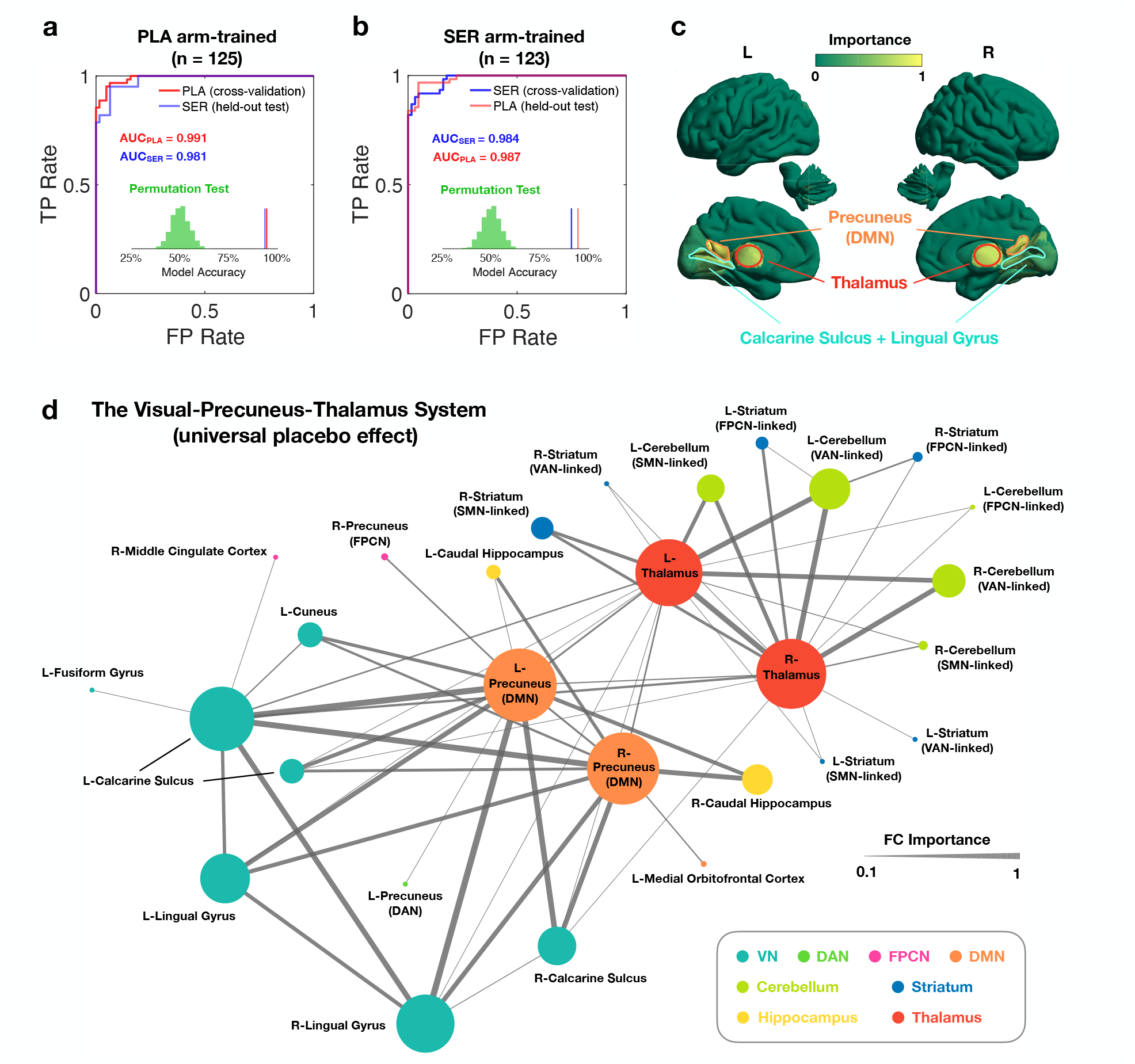
The universal one-week FC changes shared across medicated MDD patients. **a-b** Reliability and generalizability of identified biomarker. The universal treatment-induced FC changes, derived from a chronology prediction task (**Methods**), show high model accuracy (91-95%) and distinguishability (AUC) in both cross-validation and held-out test data. Cross-arm performance confirms the FC change pattern is trustable and general across treatment arms, therefore reflecting a universal placebo effect. The significance of model performance was verified by permutation tests (p < 0.001 for both arms). **c** Brain region importance in universal treatment-induced FC changes. Overall, the universal placebo effect stems from a sparse basis of brain regions. The precuneus division with dense connections to default-mode network (DMN) shows the highest importance, followed by thalamus and the visual regions involving calcarine sulcus and lingual gyrus. Both hemispheres contribute similarly with respect to these key regions. **d** Topological structure of the brain pattern for universal placebo effect. The visual-precuneus-thalamus (VPT) system, comprising 28 brain regions and 60 connections, emerges as a highly clustered connected graph with significant hemispheric symmetry. The precuneus, thalamus, and visual regions function as major hubs, with the thalamus linking to the striatum and cerebellum, the precuneus connecting to the hippocampus, frontoparietal control network (FPCN), and dorsal attention network (DAN), and regions in visual network (VN) predominantly connecting within VN and to the precuneus. Node size and line width indicate the importance of each region and connection, respectively. For **c** and **d**, the visualized brain pattern is derived from the placebo arm.

Next, we examined the key brain regions associated with the treatment-induced FC changes predictive of chronology, aiming to characterize the neural circuits underlying the universal early placebo effect. The pattern derived from the placebo arm was utilized for this analysis as it more accurately reflected the placebo effect. Overall, the universal placebo effect arose from a sparse set of brain regions (Fig. 2c). The precuneus, particularly its division with dense connections with the default-mode network (DMN), showed the strongest contribution, followed by the thalamus, calcarine sulcus, and lingual gyrus. Additionally, these regions exhibited comparably strong contributions from both hemispheres. These regions formed a highly clustered, symmetrical visual-precuneus-thalamus (VPT) system, consisting of 28 brain regions and 60 connections (Fig. 2d). Within this system, the precuneus, thalamus, and visual cortex regions served as hubs: the thalamus linked to the striatum and cerebellum, the precuneus connected with the hippocampus, frontoparietal control network, and dorsal attention network, while the visual regions primarily connected within the visual network and with the precuneus. Remarkably, all 60 FCs increased after one week of treatment, indicating enhanced functional synchronization within the VPT system (SFig. 2b).

Despite the consistent VPT change across patients, the degree of connectivity change showed significant individual-level variability (SFig. 2c). To quantify this degree of connectivity change, we calculated a VPT score for each patient, defined as the weighted sum of FC strengths dictated by the chronology prediction model. We then investigated what clinical profiles are associated with the heterogeneity in VPT score change. Foremost, the change in VPT score showed a strong negative correlation with the baseline VPT score (r = -0.780, p = 4.91 x 10^-52^, SFig. 2f), indicating reduced variability as the VPT system converges around a state with enhanced functional synchronization. Crucially, the VPT score change was significantly correlated with treatment response in the placebo arm at 8 weeks post-treatment (r = 0.193, p = 0.043, SFig. 2d), suggesting a role for VPT system early FC change in predicting degree of placebo-induced symptom alleviation. In contrast, this correlation was non-significant in the sertraline arm (r = 0.154, p = 0.124, SFig. 2e), pointing to a potential specificity of the VPT system to pure placebo effect (which may be obscured in the sertraline arm due to the combination of expectancy and medication effects present in this group). Additionally, VPT score change also significantly correlated with pre-treatment cognitive performance on the flanker task (flanker inference on accuracy: r = 0.205, p_FDR_ = 8.56 x 10^-3^, SFig. 2g), suggesting a possible link between cognitive adaptability and placebo responsiveness. No significant correlations were observed between VPT score change and other cognitive task performances, baseline MDD severity, childhood trauma history, personality traits, or mood and anxiety symptom profiles (SFig. 2g,h, SFig. 3).

**Fig. 3.**
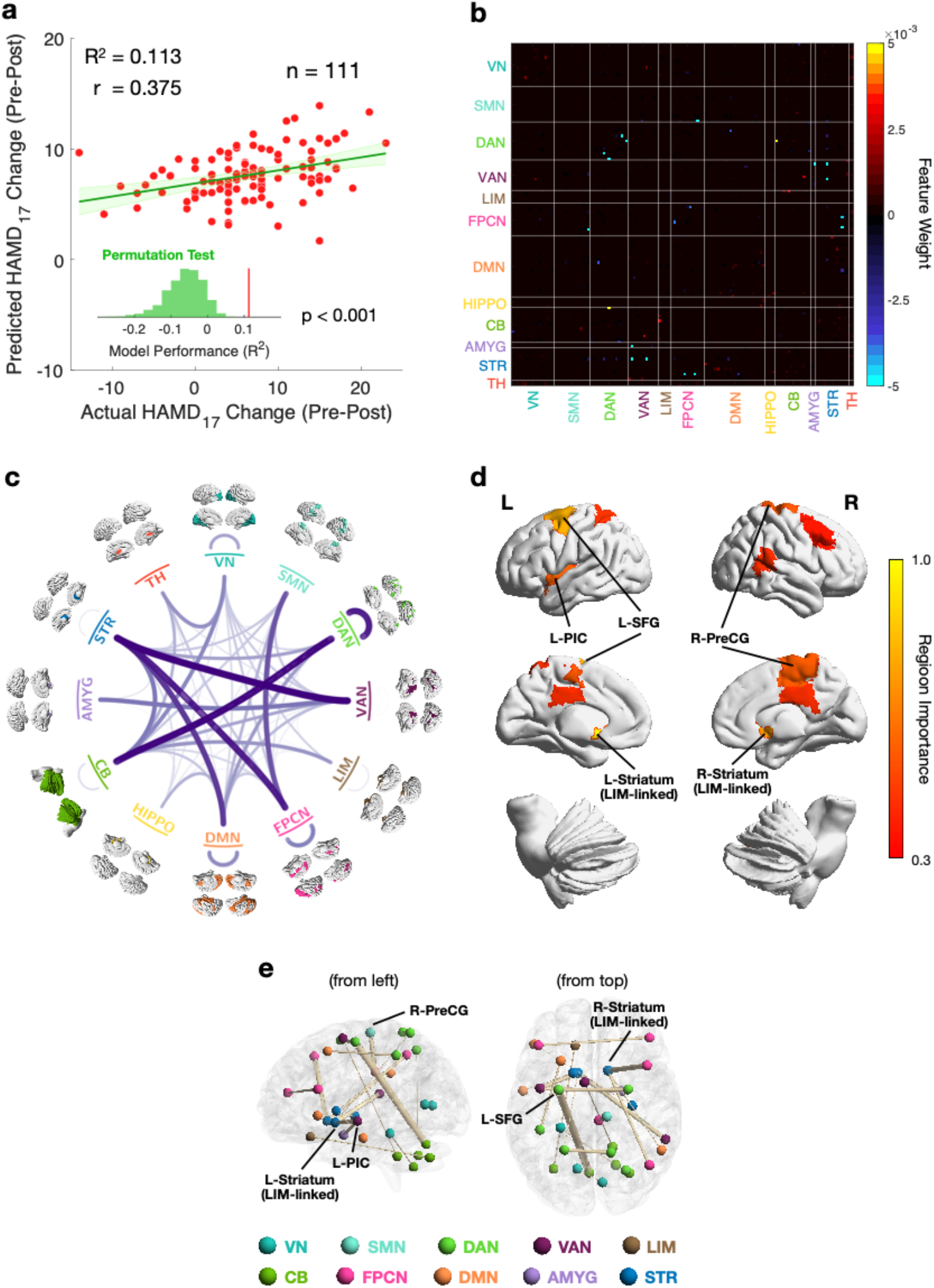
The placebo response-predictive one-week FC changes. **a** Predictability of early FC change for placebo response. **b** The FC change signature. An overall decrease in involved FCs is associated with greater placebo response. **c** Importance of network-level FC. Striatum-VAN, cerebellum-DAN, striatum-FPCN, within-DAN, and striatum-DMN connections are the most important FCs for predicting placebo response. **d** Region importance. Bilateral LIM-linked striata, left superior frontal gyrus (L-SFG), left posterior insular cortex (L-PIC), and right precentral gyrus (R-PreCG) are the brain regions with the strongest contribution to placebo response prediction. The top 10 regions are displayed for clarity. **e** Important region-level connections. The most important connections include FCs between L-SFG and the left visual-linked cerebellum, between the right LIM-linked striatum and the right middle frontal gyrus, between R-PreCG and the right midcingulate cortex, between the left LIM-linked striatum and right superior temporal gyrus, and between the left LIM-linked striatum and L-PIC. Line width indicates features weight in the prediction model. The top 20 connections are displayed for clarity.

### Response-Predictive Placebo Effect

While the VPT system change identified above reflected universal FC changes across MDD patients, it did not comprehensively capture individual variability in symptom improvement. Additionally, the modest correlation between VPT score change and treatment response in the placebo arm suggested that not all treatment-induced FC changes directly translate to clinical benefits. To better characterize the full spectrum of placebo-induced FC changes associated with symptom improvement, we developed a machine learning-based prediction model for eight-week treatment response based solely on one-week FC changes (**Methods**). This model reliably predicted placebo response in cross-validation (R^2^ = 0.113, r = 0.375), with significance confirmed by permutation testing (p < 0.001, Fig. 3a, SFig. 4). However, no significant predictability was observed for sertraline response (STable 1), suggesting that one-week FC changes primarily reflect placebo-driven mechanisms.

**Fig. 4.**
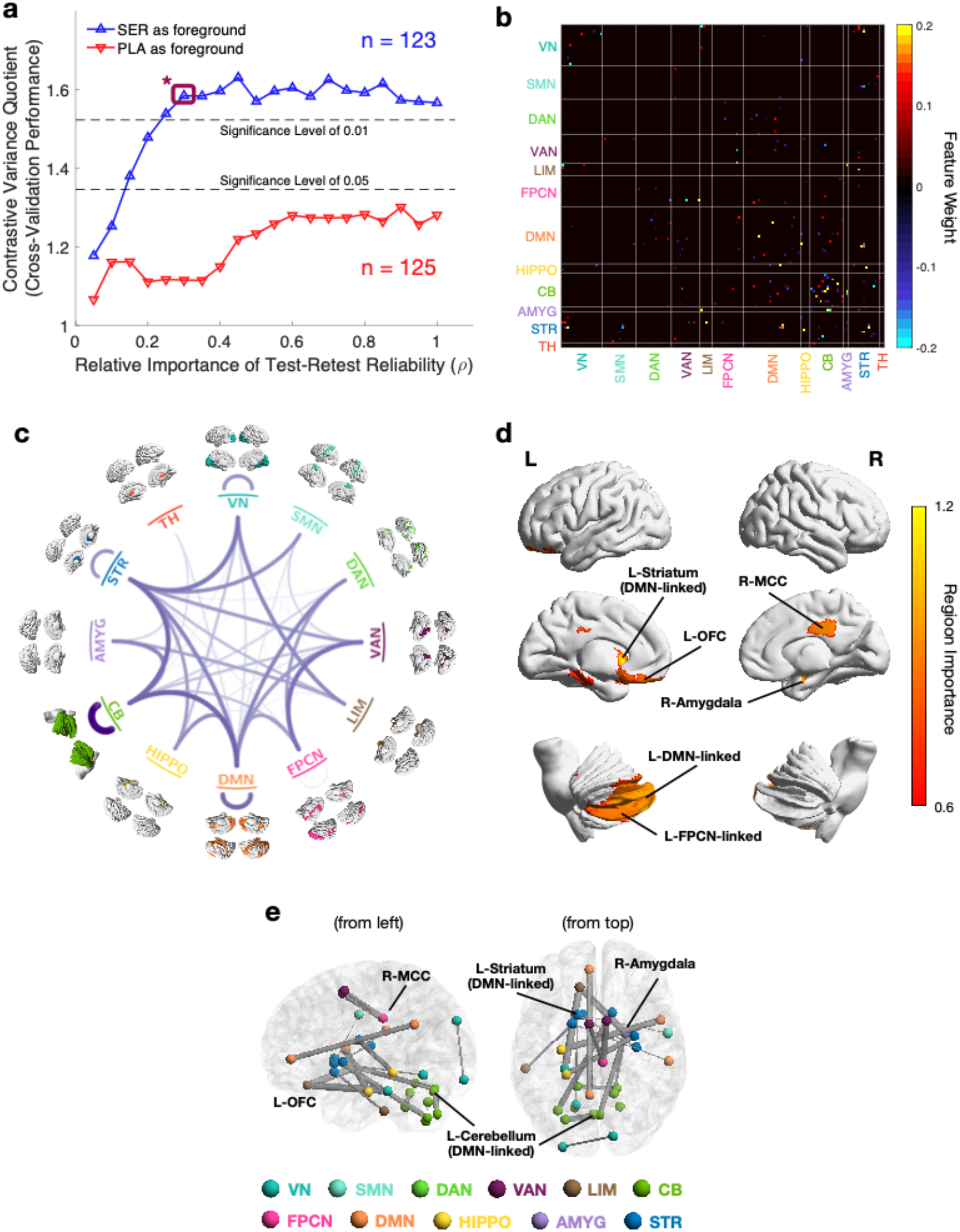
The sertraline-specific one-week FC changes representing real drug effect. **a** Significance and generalizability. The sertraline-specific FC change dimension exhibits significantly greater variance in the sertraline arm compared to the placebo arm in the validation set. In contrast, no generalizable placebo-specific dimension was identified. Introducing a test-retest reliability constraint enhances the generalizability of the drug-specific dimension, but further increasing this constraint beyond an optimal level does not improve the contrast. The operating point (brown starred box) is selected at the lowest test-retest reliability where generalizability has plateaued (*ρ* = 0.3), to maximize the sertraline-specific dimension’s distinction from placebo effects. The significance level is for the one-sided hypothesis (SER > PLA). **b** The FC change signature primarily consists of the FCs involving DMN, striatum, and cerebellum. **c** Importance of network-level FC. The real drug effect induces substantial changes in within-cerebellum connections, together with widespread connections involving DMN, striatum, and cerebellum. **d** Region importance. Left DMN-linked striatum, right amygdala, right midcingulate cortex (R-MCC), left orbitofrontal cortex (L-OFC), and the parts of left cerebellum with rich connections to DMN and FPCN are the brain regions with the strongest changes due to sertraline-specific effect. The top 10 regions are displayed for clarity. **e** Important ROI-level connections. The most important connections are FCs between L-OFC and left fusiform gyrus, between L-OFC and right amygdala, and between R-MCC and the right supplementary motor area. The top 20 connections are displayed for clarity.

Unlike the arm non-specific changes of the VPT system induced by the universal placebo effect, the brain changes predictive of greater placebo response were predominantly characterized by decreases in FC (Fig. 3b). These decreases primarily involved the striatum-VAN (ventral attention network), cerebellum-DAN (dorsal attention network), striatum-FPCN (frontoparietal control network), within-DAN, and striatum-DMN connections (Fig. 3c). Key brain regions contributing to these changes included the bilateral limbic network (LIM)-linked striata, the left superior frontal gyrus (L-SFG), the left posterior insular cortex (L-PIC), and the right precentral gyrus (R-PreCG) (Fig. 3d). The most predictive FC changes involved connections between L-SFG and the left visual-linked cerebellum, between the right LIM-linked striatum and the right middle frontal gyrus, between R-PreCG and the right midcingulate cortex, between the left LIM-linked striatum and right superior temporal gyrus, and between the left LIM-linked striatum and L-PIC (Fig. 3e).

Next, we examined the association between FC-derived response-predictive placebo effect and cognitive/clinical measures. The FC-derived placebo response prediction score correlated moderately but significantly with one-week VPT score change (r = 0.286, p = 2.33 x 10^-3^, SFig. 5a), reinforcing that overall placebo-induced FC changes partially translate to symptom improvement. The prediction score also correlated with pre-treatment choice reaction time performance (r = -0.256, p_FDR_ = 0.046, SFig. 5c), but not with other cognitive or clinical measures (SFig. 5b,c, SFig. 6), reinforcing that pre-treatment cognitive adaptability may influence placebo responsiveness. Importantly, these findings aligned with actual placebo response correlations with VPT score change (SFig. 2d) and choice reaction time (r = -0.278, p_FDR_ = 0.012, SFig. 7b), further supporting the biological and behavioral relevance of the identified striatum- and attention network-based brain signature. Beyond these associations, actual placebo response also showed significant correlations with pre-treatment HAMD_17_ score (r = 0.220, p_FDR_ = 0.027, SFig. 7a) and flanker test accuracy (r = 0.218, p_FDR_ = 0.048, SFig. 7b), however, these additional relationships were not explained by the identified brain signature. No other significant correlations were observed between placebo response and cognitive task performance or clinical measures (SFig. 7, SFig. 8).

**Fig. 5.**
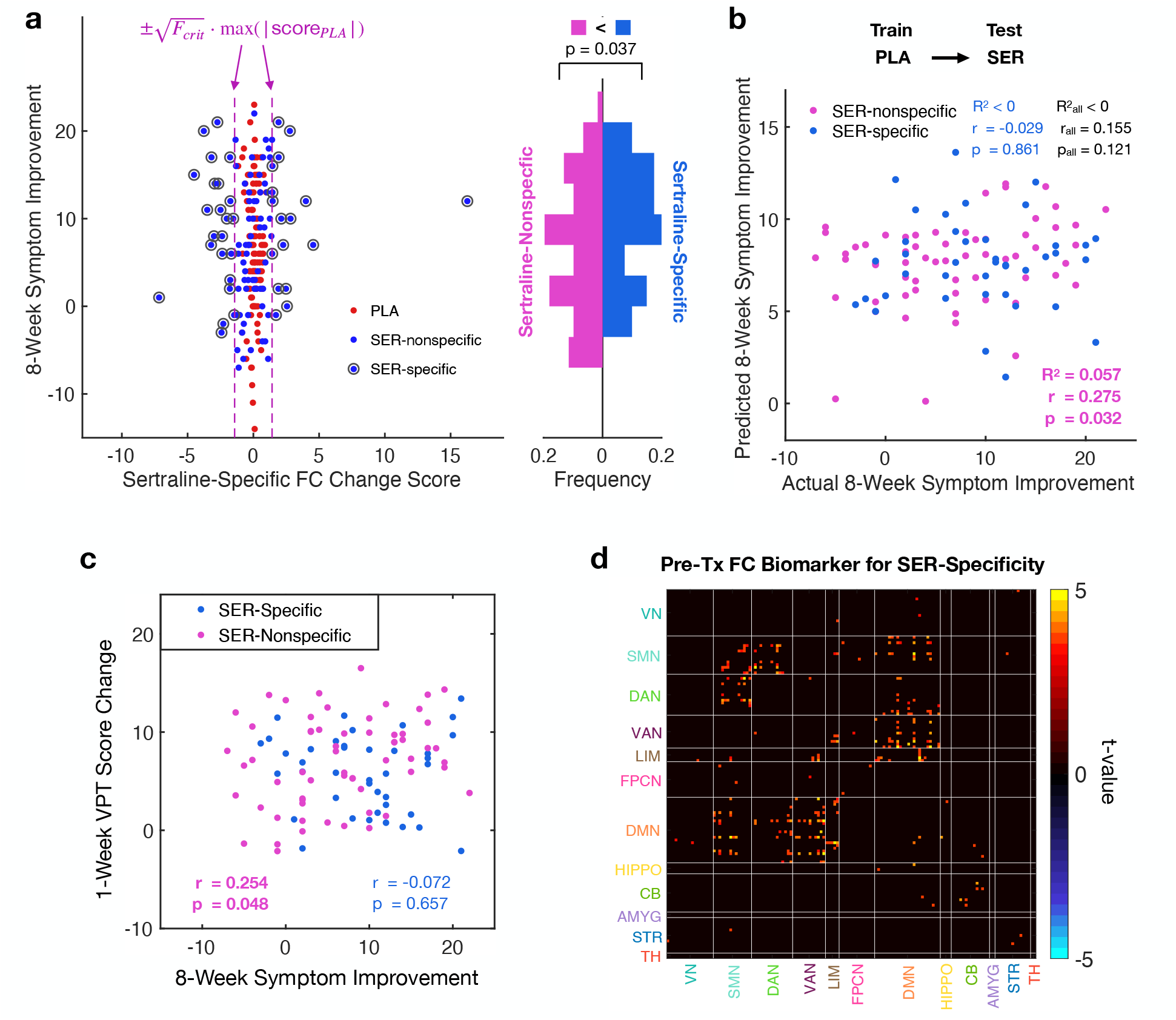
Placebo response in sertraline-medicated patients. **a** Distribution of sertraline-specific FC change scores in placebo (PLA) and sertraline (SER) arms. SER-specific FC changes are defined as absolute scores exceeding a threshold derived from the one-tailed F-test for unequal variance (SER > PLA) at p < 0.01 **(Methods**). Among 123 SER patients, 47 were classified as SER-specific and 76 as SER-nonspecific; among the 101 with available symptom improvement data, 41 were SER-specific and 61 were SER-nonspecific. SER-specific patients showed significantly greater symptom improvement. The p-value is one-tailed, based on the directional hypothesis that the efficacy of sertraline over placebo is driven by SER-specific patients. **b** Placebo-trained FC change-based prediction model for treatment response generalizes to SER-nonspecific patients. Although non-significant when applied to the full SER arm the model significantly predicts symptom improvement in SER-nonspecific patients, indicating their placebo-like response profiles. **c** Mirrored to the placebo arm (SFig. 1d), 8-week symptom improvement correlates with 1-week VPT score change in SER-nonspecific patients, supporting the placebo-like nature of their response and validating VPT change as a generalizable placebo response biomarker. **d** Pre-treatment FC profiles differentiate SER-specific from SER-nonspecific patients. The FCs involving SMN, DAN, VAN, and DMN show significant group differences, with an overall more activated state in SER-specific patients. Only features passing FDR correction are shown (*t*-test, p_FDR_ < 0.05).

**Fig. 6.**
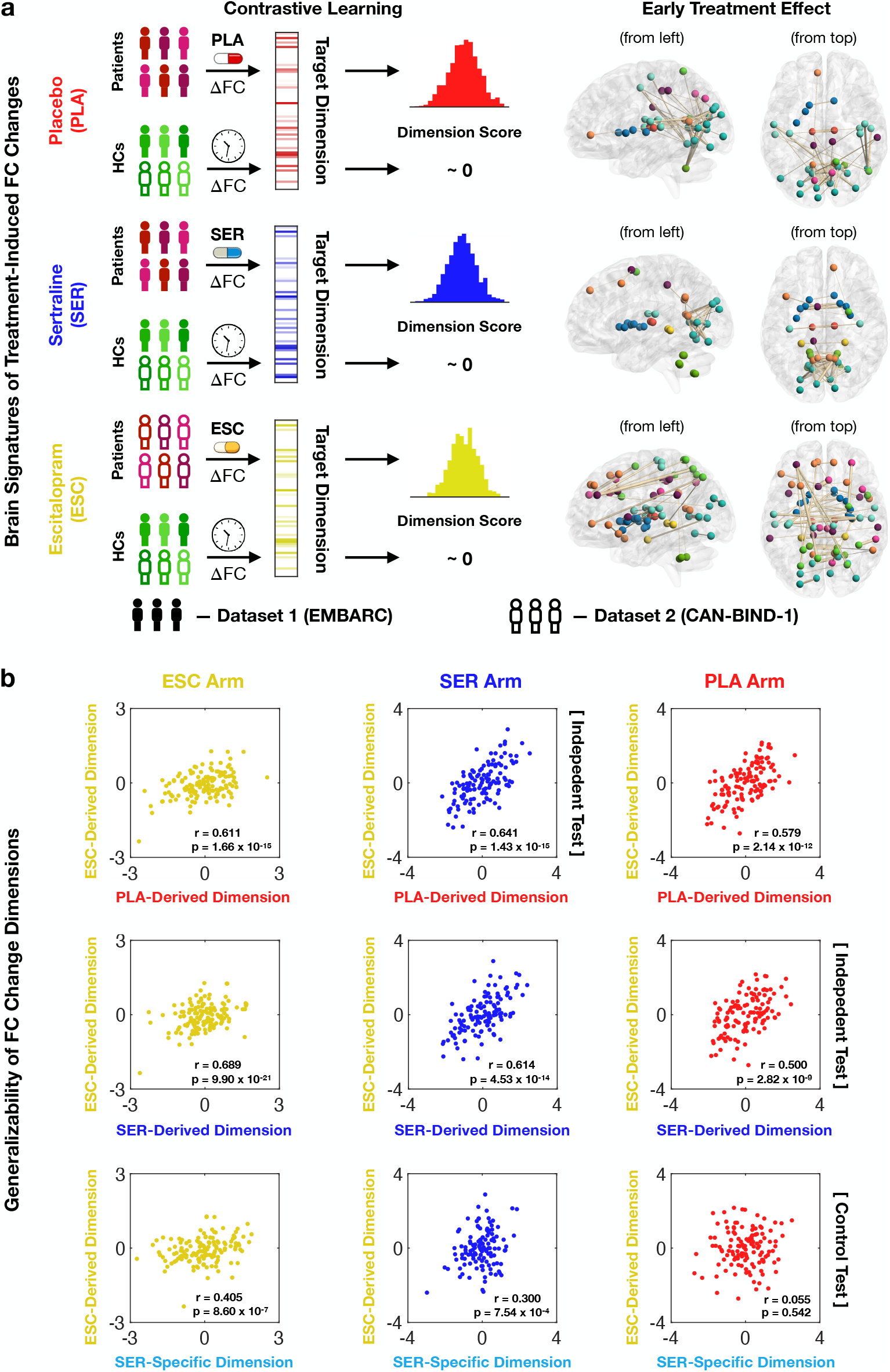
Generalizability of treatment-induced FC changes across cohorts. **a** Workflow for identifying early treatment effects and corresponding brain signatures. A contrastive learning framework is used to extract FC change dimensions that minimize variance in HCs and maximize variance in medicated-patients, separately for each treatment arm. HC data are pooled across cohorts, and their FC change — reflecting test–retest variability — is treated as a null baseline. For each region-level brain signature, the top 50 FCs are shown for clarity. **b** Generalizability of treatment-induced FC change dimensions. Cross-cohort generalizability is assessed by computing the Pearson correlation between FC change dimensions derived from the EMBARC and CAN-BIND datasets.

### Sertraline-Specific Effect

Since sertraline-medicated patients also exhibited placebo-driven FC changes, we next sought to isolate the FC changes attributable specifically to the active antidepressant. Assuming that the overall sertraline effect reflects a combination of placebo and drug-specific effects, we sought to disentangle sertraline-specific and placebo-shared FC change dimensions. To this end, we developed a contrastive variance quotient (CVQ) analysis, which identifies the dimension that maximizes the ratio of foreground and background variances (**Methods**). In this approach, FC changes from the sertraline arm were treated as foreground data, while FC changes from the placebo arm served as background data. To further distinguish true treatment effects from measurement noise, we also incorporated test-retest fMRI scans at each time point into the background, ensuring that the identified FC changes reflected genuine treatment effects rather than random variability.

Using this approach, we successfully identified a generalizable sertraline-specific FC change dimension (stratified cross-validation: p = 0.005, Fig. 4a; permutation test: p = 0.010, SFig. 9a). This drug-specific FC change dimension primarily involved cerebellum, striatum, and DMN (Fig. 4b), with particularly prominent alterations in within-cerebellum connections (Fig. 4c). Key regions contributing to these drug-specific changes included the left DMN-linked striatum, the right amygdala, the right midcingulate cortex (R-MCC), the left orbitofrontal cortex (L-OFC), and the left cerebellar hemisphere with strong connections to DMN and FPCN (Fig. 4d). The most sertraline-specific changes involved the FCs between L-OFC and left fusiform gyrus, between L-OFC and right amygdala, and between R-MCC and the right supplementary motor area (Fig. 4e).

Notably, incorporating test-retest reliability terms significantly improved the generalizability of the identified FC change dimension (Fig. 4a, SFig. 9b), underscoring the importance of controlling for run-to-run fluctuations. To confirm that the identified drug-specific changes were not false positives, we conducted the same analysis using the placebo arm as the foreground data and the sertraline arm as the background. No generalizable placebo-specific FC changes were identified using the placebo arm as foreground (Fig. 4a), supporting the premise that the overall sertraline effect reflects both placebo and real drug effects, whereas the placebo effect is purely placebo-driven.

### Placebo Response in Sertraline Arm

While we identified generalizable sertraline-specific FC changes, further investigation revealed that not all sertraline-medicated patients exhibited these drug-specific patterns (Fig. 5a). This finding echoed with our observation based on predictive modeling that one-week FC changes could not reliably distinguish between sertraline and placebo treatments (STable 2), collectively suggesting that early FC changes in some sertraline-medicated patients may be indistinguishable from placebo effects. To explore this further, we empirically categorized sertraline-treated patients into two subgroups based on the presence or absence of significant sertraline-specific FC changes (**Methods**): the sertraline-specific and sertraline-nonspecific groups. Approximately 40% of sertraline-medicated patients were classified as sertraline-specific (n = 47), while the remaining patients showed no significantly greater FC changes than placebo-treated patients and were classified as sertraline-nonspecific (n = 76).

Given that sertraline-nonspecific patients exhibited only placebo-like FC changes, we hypothesized that their symptom improvement was primarily driven by placebo effects. To test this, we evaluated whether the placebo response prediction model could generalize to the sertraline-nonspecific group. Remarkably, while the model failed to predict overall sertraline response (R^2^ < 0, r = 0.155, p = 0.121), it significantly predicted symptom improvement in sertraline-nonspecific patients (R^2^ = 0.057, r = 0.275, p = 0.032, Fig. 5b), supporting the hypothesis that their response was placebo-driven. Furthermore, while symptom improvement in the full sertraline arm did not correlate with VPT score changes (SFig. 2e), a significant correlation emerged within the sertraline-nonspecific subgroup r = 0.254, p = 0.048, Fig. 5c). This mirrored the placebo arm results (SFig. 2d), reinforcing that sertraline-nonspecific patients responded as if treated with placebo and validating the VPT score change as a generalizable biomarker of placebo response.

Next, we examined whether this subgrouping could explain the modest but significant superiority of sertraline over placebo. Encouragingly, sertraline-specific patients exhibited significantly greater symptom improvement than sertraline-nonspecific patients (one-tailed t-test: p = 0.037, Fig. 5a). Since the improvement in sertraline-nonspecific patients was primarily placebo-driven, this finding suggests that the higher efficacy of sertraline is largely attributable to sertraline-specific patients, highlighting the clinical importance of identifying sertraline-specific responders. To improve its clinical applicability, we investigated the pre-treatment FC signatures that could differentiate these subgroups. We found that FCs involving brain regions in the somatomotor network (SMN), DAN, VAN, and DMN significantly differed between subgroups (t-test: p_FDR_ < 0.05), with sertraline-specific patients showing an overall highly-connected state (Fig. 5d, SFig. 10a-c). Notably, this somatomotor-attention-default-mode dimension formed a unified system involving FC features highly intercorrelated with each other (SFig. 10d). We also developed a pre-treatment FC-based classification model to distinguish these subgroups, providing a brain signature that may serve as a biomarker to identify sertraline-specific patients at baseline (SFig. 11). No pre-treatment structural connectivity features and clinical measures distinguished these subgroups (SFig. 12, SFig. 13).

### Generalizability on Independent Cohort

Generalizability is crucial for the clinical translatability of neuroimaging biomarkers. To evaluate the generalizability of our identified treatment-induced FC changes, we used a second dataset consisting of MDD patients treated with escitalopram, with fMRI data scanned at baseline and two weeks after treatment onset. Since the two datasets differed in fMRI acquisition timing and medication type, we focused on evaluating the replicability of general treatment-induced FC changes and did not tailor the brain dimensions to any clinical targets. Specifically, we applied CVQ analysis using FC changes from untreated healthy controls (HCs) and test-retest fluctuations across subjects and time points as background data to each patient group: placebo, sertraline, and escitalopram. This allowed us to identify overall treatment effects of each medication, with escitalopram-induced FC changes from the replication dataset serving as a reference for evaluating the generalizability of FC changes derived from the discovery dataset (Fig. 6a, **Methods**).

As a result, we obtained three FC change signatures representing the placebo, sertraline, and escitalopram effects (Fig. 6a), all of which showed significant cross-validation performance (SFig. 14). We first compared the placebo- and escitalopram-derived FC change dimensions using the sertraline arm as an independent cohort, as it was not used in generating these dimensions. Remarkably, the dimension scores of placebo- and escitalopram-derived FC changes were strongly correlated across sertraline-treated patients (r = 0.641, p = 1.43 x 10^-15^, Fig. 6b), demonstrating high replicability of treatment-induced FC changes across cohorts and suggesting that placebo effect also significantly contributes to escitalopram’s overall treatment effect. Similar strong correlations were observed in the placebo and escitalopram arms (placebo: r = 0.579, p = 2.14 x 10^-12^; escitalopram: r = 0.611, p = 1.66 x 10^-15^, Fig. 6b). Next, we compared the sertraline- and escitalopram-derived FC change dimensions using the placebo arm as an independent cohort. Again, the dimension scores of sertraline- and escitalopram-derived FC changes exhibited strong correlations in the placebo arm (r = 0.500, p = 2.82 x 10^-9^, Fig. 6b) and were similarly correlated in the sertraline and escitalopram arms (sertraline: r = 0.614, p = 1.43 x 10^-15^; escitalopram: r = 0.689, p = 9.90 x 10^-21^, Fig. 6b). The correlation in the placebo arm reflects the generalizability of the placebo effect in both sertraline- and escitalopram-induced FC changes, reinforcing that treatment-induced FC changes are generalizable across cohorts and medications. However, while the strong correlation between sertraline- and escitalopram-derived FC change signatures suggests overall similarity, this alone does not confirm shared real drug effects, as the placebo effect itself could account for the overlap. To address this, we compared the sertraline-specific FC change dimension with escitalopram-derived FC change dimension. We found significant correlations across patients in both the sertraline and escitalopram arms (sertraline: r = 0.300, p = 7.54 x 10^-4^; escitalopram: r = 0.405, p = 8.60 x 10^-7^, Fig, 6b), confirming a shared and generalizable real drug effect between these two antidepressants. Notably, the correlations involving the sertraline-specific dimension were weaker than those observed for overall treatment effects. This finding is consistent with our earlier observation that one-week sertraline-induced FC changes are largely dominated by the placebo effect, and suggests two-week escitalopram-induced changes may follow a similar pattern. Importantly, the sertraline-specific and escitalopram-derived dimension scores were not significantly correlated in the placebo arm (r = 0.055, p = 0.542, Fig. 6b), confirming that sertraline-specific FC changes were absent in placebo-treated patients.

## Discussion

In this study, we systematically characterized early FC changes induced by antidepressants and placebo. We focused on three major facets: 1) the universal placebo effect shared across medications and levels of treatment responsiveness, 2) the placebo effect predictive of symptom improvement, and 3) the drug-specific effect detectable only in patients receiving active antidepressants. Remarkably, with evidence spanning from brain changes to symptom improvement, we identified a subgroup of sertraline-treated patients whose early neurobiological responses to treatment resembled those receiving placebo. We further showed that the modest overall superiority of sertraline over placebo may be driven by greater symptom improvement in patients displaying prominent early sertraline-specific neurobiological change patterns compared to those not displaying those patterns despite receiving the same treatment. Meanwhile, the generalizability of placebo-derived results on the sertraline-nonspecific subgroup supports the reliability of this placebo signature and converges with clinical intuition that patients less responsive to antidepressant treatment may still benefit, to a lesser extent, from the placebo response component. These early FC changes were further validated using an independent cohort treated with escitalopram, confirming that treatment-induced FC changes are generalizable across the SSRI class of medications and across patient cohorts. Together, our findings illustrate a novel conceptual and analytical framework for parsing apart different components of treatment-related changes, mark a substantial advance in understanding the neurobiological underpinnings of antidepressant treatment, and provide new opportunities for early treatment optimization in MDD clinical care.

Previous research has identified post-treatment FC changes associated with symptom improvement^12^, yet it remains unclear whether non-responders fail to show any FC alterations or if their brain changes simply do not translate into behavioral changes. Our results suggest the latter: treatment induces widespread early FC changes, even in the absence of symptom improvement. Consistent across all medicated patients, this universal placebo effect manifests as increased FC within a VPT system comprising the visual network, precuneus, and thalamus. Interestingly, these brain regions are also the ones with the densest local connections and the highest level of concurrent metabolic need and activity^18^, implicating a potential link between placebo-related FC changes and cerebral metabolism. The observed correlation between the magnitude of VPT FC increase and symptom improvement further supports this interpretation, aligning with prior work suggesting a relationship between antidepressant response and metabolic pathways^19, 20^. These findings position the early universal placebo effect as a clinically relevant biomarker in the pathway linking primary treatment-induced FC changes to symptom relief.

We next examined the placebo-induced FC changes predictive of symptom improvement, which prominently featured decreases in FCs involving the striatum and attention networks. This finding aligns with prior evidence that post-treatment FC changes in the striatum are associated with treatment responsiveness^21^, suggesting these striatal alterations may emerge within one week of treatment. This interpretation is further supported by the observation of increased endogenous opioid release in the nucleus accumbens following one week of placebo administration^7^, indicating early neurotransmission changes in striatal circuitry. In contrast to pre-treatment biomarkers of antidepressant response^5, 22-26^ and MDD symptomatology^25-29^, the anterior cingulate cortex, posterior cingulate cortex, and prefrontal cortex did not emerge as important brain regions in the early FC placebo change signature for symptom improvement. This observation is consistent with prior metabolic imaging studies showing that placebo-induced metabolic changes in these cortical regions emerge after six weeks of treatment, but are not yet detectable at one week^6^. Collectively, these results reinforce the biological plausibility of our early FC biomarker, linking it to both metabolic and neurotransmission processes. They also highlight the unique mechanistic insight provided by capturing treatment-induced FC changes at early time points, offering a new perspective on how initial neural adaptations may mediate subsequent clinical improvement.

We also delineated the drug-specific effects associated with sertraline treatment. Notably, the sertraline-specific FC changes centered on the left cerebellum, particularly in regions functionally connected to the DMN and FPCN, consistent with prior evidence that SSRIs can induce acute FC alterations in cerebello-cortical circuits involving these networks^30^. Aligning with earlier findings^31, 32^, sertraline also induced FC changes in the striatum and amygdala. These brain regions have been independently implicated in MDD pathophysiology and treatment response, with previous studies identifying regional homogeneity in the left cerebellum^33^, FC between cerebellum and DMN/FPCN^34^, FC between striatum and DMN^35^, and amygdala activity level^32^ as biomarkers for MDD diagnosis or antidepressant response. These converging lines of evidence may explain the superior symptom improvement observed in patients exhibiting sertraline-specific FC changes. Our findings thus support the clinical utility of drug-specificity subgroups: pre-treatment FC features may aid in identifying patients more likely to benefit from sertraline, while early FC changes can inform dynamic treatment adjustment, ultimately contributing to more personalized and effective interventions for MDD.

In contrast to drug-specific FC changes derived from group-level comparison^12^, the sertraline-specific dimension we identified captures the variability of individual-level FC changes – the magnitude and direction of drug-induced FC changes can differ across patients. This approach allowed us to parse the heterogeneity of sertraline’s effect and revealed that a subset of sertraline-medicated patients exhibit placebo-like FC changes and symptomatic response. While prior studies have interpreted the lack of generalizability between sertraline and placebo response prediction models as evidence for biomarker specificity^36-38^, this interpretation may overlook the heterogeneity within the sertraline arm and underestimate the influence of placebo effect. Our findings instead show that the placebo response model is indeed generalizable -- to the subgroup of sertraline-medicated patients lacking drug-specific FC changes. Combined with the generalizability of the placebo-specific correlation between VPT score change and treatment outcome, these results strongly support that a subset of antidepressant-medicated patients symptomatically improve primarily through placebo mechanisms, and these individuals tend to show less dramatic symptom improvements. Beyond the implications for personalized treatment, these findings illustrate a more nuanced understanding of antidepressant response: drug response is not always attributable to the active antidepressants -- the placebo effect is both real and stable^39^ and, for some patients, may be the primary driver of clinical benefit.

Remarkably, we demonstrated that early FC changes identified in the EMBARC cohort generalized to the CAN-BIND-1 cohort. Although direct replication of prediction tasks was not feasible due to differences in medication type and fMRI acquisition timing, we leveraged HCs to isolate general treatment effects from random effects and test-retest variability. This approach revealed that placebo-induced FC changes are generalizable across cohorts and medication arms, and that a shared drug-specific effect exists between sertraline and escitalopram -- consistent with the well-established reproducibility of the placebo effect^39^ and the common pharmacological mechanism of SSRIs^40^. Notably, we highlight the methodological innovation -- incorporating SNR into the learning objective -- that may contribute to the exceptional replicability of our findings. This constraint allowed us to isolate meaningful treatment-related FC changes from random fluctuations, improving data quality and replicability. By organizing FC features into coherent, interpretable brain dimensions, our approach also enhanced signal synergy and interpretability. We recommend that future resting-state fMRI studies adopt SNR-aware objectives to improve the reliability and reproducibility of neuroimaging-based biomarkers.

Despite the comprehensive scope of our study, several important questions remain. First, we did not identify an early FC change signature that reliably predicts individual-level sertraline response. This may reflect the dominance of placebo effect at early stages of treatment, highlighting the need for future work mapping the time course of drug-specific FC changes that are predictive of treatment response. Additional measurement timepoints further into the treatment course would also resolve the important question of whether the early sertraline-specific changes detected here may appear later in some patients found to lack such changes early in the treatment course, which would address whether the early temporal emergence of these changes is crucial as opposed to their overall presence or absence. Second, due to the absence of a placebo arm in the CAN-BIND-1 cohort, we were unable to determine whether sertraline and escitalopram exhibit distinct early drug-specific FC changes. We advocate for future studies to include a placebo control or a placebo lead-in period prior to active antidepressant administration to better disentangle actions of antidepressants from placebo effect. Further research is also needed to elucidate the neural circuits mediating the translation of early FC changes into symptom improvement. Finally, clinical trials are warranted to test the utility of drug-specific subgrouping for guiding personalized antidepressant treatment and improving outcomes in MDD.

In summary, we systematically characterized early FC changes induced by antidepressants and placebo, uncovering three distinct facets: a universal placebo effect, a response-predictive placebo effect, and a drug-specific effect. Notably, these FC changes were generalizable across independent cohorts and medication types. We found that early FC changes emerge in nearly all medicated patients -- regardless of medication type or clinical response -- yet only a subset of these changes contribute to symptom improvement. While early FC changes were largely dominated by placebo effect, we also identified drug-specific alterations that were associated with treatment outcome. Based on these findings, we proposed a biologically informed subgrouping criterion grounded in the presence or absence of drug-specific FC changes. Patients lacking drug-specific changes responded similarly to placebo-treated patients, suggesting that the observed superiority of sertraline over placebo may reflect underlying patient-specific drug responsiveness. Collectively, our findings fill a critical knowledge gap regarding early treatment-induced FC changes in MDD, offer new insights into the neural underpinnings of placebo and pharmacological effects, and establish a foundation for future precision medicine approaches leveraging early brain changes to guide adaptive treatment.

## Methods Participants

### Discovery Dataset – The EMBARC Cohort

The <u>E</u>stablishing <u>M</u>oderators and <u>B</u>iosignatures of <u>A</u>ntidepressant <u>R</u>esponse in <u>C</u>linical Care (EMBARC) study recruited 296 MDD patients and 52 HCs aged between 18 and 65, who were randomly assigned to two treatment arms receiving either sertraline or placebo^41^. Treatment randomization was stratified to ensure balanced distributions of study site, depression severity, and chronicity across treatment arms. Eligible patients were required to have MDD as their primary diagnosis by the Structured Clinical Interview for Diagnostic and Statistical Manual of Mental Disorders, Fourth Edition (DSM-IV) Axis I Disorders^42^, a Quick Inventory of Depressive Symptomatology score ≥ 14, and an onset of MDD episode prior to age 30. Additionally, recruited patients must not have experienced failure with any antidepressant during the current episode. Key exclusion criteria included: 1) ongoing pregnancy or breastfeeding, 2) sexual activity without contraception, 3) a lifetime history of psychosis or bipolar disorder, 4) substance dependence in the past six months or substance abuse in the past two months, 5) psychiatric or general medical conditions requiring hospitalization, 6) contraindication to sertraline or bupropion, 7) clinically significant laboratory abnormalities, 8) a history of epilepsy or other conditions requiring anticonvulsants, 9) electroconvulsive therapy, vagal nerve stimulation, transcranial magnetic stimulation or other somatic treatments in the current episode, 10) ongoing medication intake (antipsychotics and mood stabilizers, etc.), 11) ongoing psychotherapy, and 12) significant suicide risk. The EMBARC study adhered to FDA guidelines and the Declaration of Helsinki and was approved by the Institutional Review Board of each clinical site. All participants have signed informed consent and given agreement to all study procedures before enrollment.

With 9 patients dropped out before receiving the first dose and 22 discontinued during the treatment, 137 patients in the sertraline arm and 142 in the placebo arm completed one week of treatment, among which 129 sertraline-treated and 136 placebo-treated patients completed the eight-week full course of treatment (SFig. 15). Medication dosing began at 50mg and was gradually increased to a maximum of 200 mg, based on patient responsiveness and tolerance. Treatment response was measured as the change in the 17-item Hamilton Depression Rating Scale (HAMD_17_) score from pre-to post-treatment. Patients without available endpoint HAMD_17_ scores were excluded from the study. fMRI scans were collected for MDD patients both before the treatment and 1 week after treatment onset. For feature quality, only the subjects with two high-quality resting-state fMRI runs at both time points were retained, resulting in 123 participants in the sertraline arm, 125 in the placebo arm, and 29 HCs (SFig. 15). Each patient may have one or two fMRI recordings at each time point. Diffusion tensor imaging (DTI) scans were collected at baseline (see **Supplementary Methods** for DTI acquisition and processing).

### Replication Dataset – The CAN-BIND-1 Cohort

The initial samples of <u>Can</u>adian <u>B</u>iomarker Integration <u>N</u>etwork in <u>D</u>epression (CAN-BIND)-1 study included 196 MDD patients and 110 HCs^43^. The inclusion criteria^44^ were: 1) outpatients aged 18 to 60, 2) experiencing a major depressive episode diagnosed by DSM-IV Text Revision and confirmed by the Mini International Neuropsychiatric Interview, 3) depressive episode duration ≥ 3 months, 4) Montgomery-Åsberg Depression Rating Scale (MADRS) score ≥ 24, and 5) sufficient English proficiency to complete interviews and self-report questionnaires. The exclusion criteria included: 1) bipolar I or II disorder, 2) primary diagnosis of psychiatric disorders other than MDD, 3) significant personality disorder that might interfere with study participation (e.g., borderline, antisocial), 4) high suicidal risk, 5) substance dependence or abuse in the past six months, 6) significant neurological disorders, head trauma, or other unstable medical conditions, 7) ongoing pregnancy or breastfeeding, 8) psychosis in the current episode, 9) high risk for hypomanic switch, 10) resistance to at least four antidepressant medications at therapeutic dosages, 11) previous unresponsiveness or contraindication to escitalopram or aripiprazole, 12) psychological treatment within past three months with intent to continue, and 13) contraindications to MRI. The CAN-BIND-1 study was approved by the ethics committees of each participating clinical site. All eligible participants have provided written informed consent for all study procedures at the screening visit.

For patients who had taken psychoactive medications, a wash-out period of at least five half-lives of medication was required. Subsequently, all MDD patients were treated with 10-20 mg/day escitalopram for eight weeks. fMRI scans were performed at baseline and two weeks after treatment onset. 138 MDD patients and 87 HCs have two fMRI runs at both time points.

### Functional MRI Acquisition

In the EMBARC study, resting-state fMRI data were collected using a single-shot gradient echo-planar pulse sequence. While different scanners were used at four clinical sites (Columbia University: General Electric 3T; Massachusetts General Hospital: Siemens 3T; University of Texas Southwestern Medical Center: Philips 3T; University of Michigan: Philips 3T), acquisition parameters were standardized. Key parameters included: 2000 msec repetition time, 28 msec echo time, 90° flip angle, 64 × 64 matrix size, 3.2 × 3.2 x 3.1 mm^3^ voxel size, 39 axial slices, and 180 image volumes.

In the CAN-BIND-1 study, resting-state fMRI data were acquired over a 10-minute scan using a whole-brain T2*-sensitive blood-oxygen-level-dependent (BOLD) echo planar imaging sequence. Participants were instructed to remain still with their eyes open, focusing on a fixation cross^45^ throughout the scan. Key acquisition parameters, consistent across study sites, included a repetition time of 2000 msec, an echo time of 30 msec, and voxel dimensions of 4 mm × 4 mm × 4 mm^46^. Scanners varied by site (GE 3T for Toronto Western/Toronto General Hospital, Centre for Addiction and Mental Health, McMaster University, and University of Calgary; Philips 3T for University of British Columbia; Siemens 3T for Queen’s University), with minor protocol variations^46^.

### Functional MRI Preprocessing

Resting-state fMRI data were preprocessed using the fMRIPrep pipeline^47^. First, T1-weighted images were corrected for intensity nonuniformity and underwent skull stripping. Spatial normalization was conducted via nonlinear registration with the T1-weighted reference^48^. Brain tissue was segmented from the reference skull-stripped T1-weighted image^49^. Fieldmap data were used to correct low- and high-frequency distortions due to field inhomogeneity, enabling a more accurate co-registration with anatomical reference and an echo-planar imaging reference to be obtained with these distortions corrected. The BOLD reference was then transformed to the T1-weighted image using boundary-based registration^50^. Afterward, a 6 mm full-width half-maximum isotropic Gaussian kernel was employed to remove non-steady-state volumes and smooth spatial signals, followed by an independent component analysis^51^ to automatically remove motion artifacts from the preprocessed BOLD time-series. Lastly, quality control excluded data with significant head motion (>0.5 mm framewise displacement or BOLD signal displacements >0.5%)^52, 53^.

### Functional Connectivity Calculation

The preprocessed fMRI data were first averaged into time series for 135 brain regions, based on the Schaefer atlas of 100 parcels^54^ and 35 subcortical parcellations including striatum, cerebellum, amygdala, hippocampus, and thalamus^55, 56^. Subsequently, FC was calculated as the Pearson’s correlation coefficient between time series of each region pair. Lastly, Fisher’s r-to-z transformation was applied to the FC features to improve normality.

### Treatment-Induced FC Change-Based Prediction

Our primary aim is to investigate the FC changes induced by antidepressant treatment and placebo. By incorporating brain dimension and signal-to-noise ratio (SNR) into a regularized end-to-end learning procedure, we developed an innovative framework that, 1) Accounts for the synergy across FC features, 2) Differentiates treatment-induced FC changes from random fluctuations, and 3) Specifically examines the association between treatment-induced FC changes and targets of interest. The overall learning objective is formularized as a combination of SNR constraint, prediction task, and regularization:

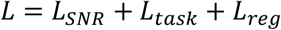

As detailed below, our proposed framework yields robust treatment-induced FC biomarkers with high interpretability and replicability. To ensure fair regularization, FC features need to be normalized across subjects within each cross-validation fold to have zero mean and unit variance. Notably, this normalization operation is conducted with both time points pooled together. Otherwise, the difference in mean across time points, which is essentially the treatment-induced change we seek, would be eliminated as means are set to equal. The functional connectivity matrix and prediction target are also scaled to have unit Frobenius norm to enhance the stability of optimization procedure.

### Brain Dimensions

Our previous study demonstrated that integrating latent dimension identification into prediction frameworks enhances the interpretability and stability of predictive patterns^4^. Inspired by this work, we introduce the concept of brain dimension to improve feature reliability and distinguish between treatment-induced FC changes and its natural variability. Conceptually, a brain dimension represents an FC community whose members collectively achieve a specific function. Given an FC data matrix *X* and brain dimension composition *W*, the dimension score is 𝒢 = *XW*. Remarkably, the importance of FC features may vary within a dimension, making each dimension composition a weighted sum rather than a binary membership.

Given their essence, we propose two key assumptions for brain dimensions in our framework. 1) <u>Consistent composition</u>. Although interactions between brain regions may change during treatment, the function of each brain region remains largely consistent. Thus, the composition of each brain dimension, associated with a latent function, remains stable over time. This means the brain dimension composition at baseline (week 0) and week 1 should be identical: *W* = *W*(*t* = 0) = *W*(*t* = 1). Notably, the dimension scores at baseline 𝒢_0_ = 𝒢(*t* = 0) and one week after treatment onset 𝒢_1_= 𝒢(*t* = 1) may differ, with the treatment-induced change in dimension score (𝒢_1_− 𝒢_0_) being of primary interest. 2) <u>Test-retest reliability</u>. The identified brain dimension scores should be reliable metrics. Therefore, dimension scores from different fMRI runs collected at the same time point for the same subject should agree. Denoting the data matrix for week i’s j-th fMRI run as *X*_*ij*,_ this implies *X*_01,_*W* ≈ *X*_02_*W* and *X*_11_*W* ≈ *X*_12_*W*. This test-retest reliability constraint is implemented via the SNR loss. Also, treatment-induced brain dimension score change serves as the features for prediction tasks, reducing the feature dimensionality and alleviating overfitting issues.

### Maximized SNR

Resting-state fMRI-based FC is known to have a low SNR^15^ and high test-retest variability^16^ due to various noise sources, limiting the replicability of its findings^17^. This unfavorable observation poses a particular challenge for FC change-based prediction frameworks, as FC changes are especially sensitive to test-retest reliability. To address this, we introduce an SNR loss to the overall learning objective to distinguish FC changes induced by treatment against its natural variability.

Essentially, we seek more robust estimates of dimension scores given two fMRI runs at each time point. As the reliability and noise power depend on the deviations of the dimension scores derived with each fMRI run from the overall estimate, the noise power can be formularized as:

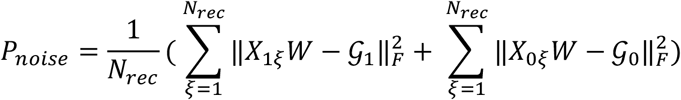

where *N*_*rec*_ = 2 denotes the number of fMRI runs at each time point for each subject, and 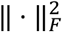 is the Frobenius norm. Minimizing this noise power is essentially minimizing the difference between *X*_01_ *W* and *X*_02_ *W*, and *X*_11_ *W* and *X*_12_ *W*. In contrast, signal power reflects the magnitude of differences in dimension scores across time points, which can be quantified by the deviations of the dimension scores derived with each fMRI run from the grand mean of all runs:

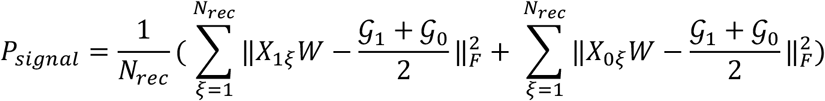

To maximize SNR, we define SNR loss as the inverse of 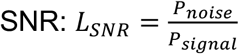. Notably, this SNR loss is closely related to the coefficient of determination (R^2^) – signal power represents the total sum of squares and noise power the residual sum of squares. Therefore, while established for two recordings and two time points, this SNR loss is readily generalizable to cases with more recordings and time points.

### Prediction Task

The prediction loss in our learning objective is based on brain dimensions. To investigate treatment-induced FC changes, we focus on target prediction using brain dimension score changes as input features. For regression tasks, the prediction loss is formularized as:

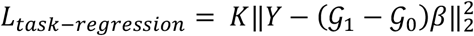

where *K* is a hyperparameter controlling the relative importance of the prediction task in the overall learning objective, *Y* is the prediction target (e.g., treatment response), and *β* represents the model weight of predictive features. For binary classification tasks, the prediction loss is:

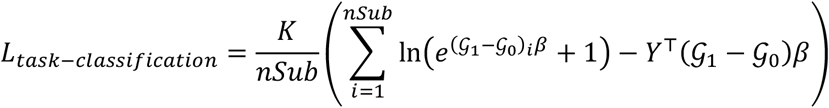

where *nSub* is the number of subjects. *Y*_*i*_ ∈ {0,1} is the binary label of prediction target (e.g., treatment arm). For multiclass cases, *L*_*task*_ can be formularized with the softmax function.

Crucially, brain dimension identification and the prediction task form an integrated process, instead of being separated into different stages. They are simultaneously conducted, allowing for a collective adjustment of brain dimensions and the predictive pattern throughout optimization, with the reliability of updated brain dimensions ensured by the SNR loss (SFig. 16).

### L0-regularization on Brain Dimensions and Predictive Pattern

Our prior work showed that L0-regularization effectively identifies distinct target-predictive dimensions and reduces their cardinality, two feats unattainable by L1-regularization^4^. Therefore, we continue using L0-regularization to achieve sparsity in the loading of brain dimensions and predictive patterns, enhancing the interpretability and generalizability of identified biomarkers. The regularization term in the loss function is formularized as:

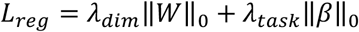

where *λ*_*dim*_ and *λ*_*task*_ are sparsity hyperparameters for brain dimensions and the model weight of prediction task. We employed the continuous sparsification technique^57^ to optimize the L0-regularized loss function. For details about the optimization procedure, see **Appendix A**.

### Chronology Prediction Task for Universal Placebo Effect Identification

Compared with correlation analyses or statistical tests conducted on all samples, prediction frameworks offer more generalizable results by leveraging cross-validation and held-out data testing. Therefore, we propose a chronology prediction task to identify more replicable brain patterns associated with treatment-induced FC changes (SFig. 1).

Our goal is to derive a biomarker that can distinguish whether an fMRI recording is from the pre-treatment session or one week after treatment onset. Given the heterogeneous baseline conditions of MDD patients, our primary aim is to differentiate fMRI recordings from the two time points for the same subject rather than pooling baseline and week 1 data as two classes. This approach also allows us to identify the general treatment effect that is universal at the individual level.

In this design, a pair of fMRI recordings at two time points for the same patient form a single data point, with the chronology label (chronological vs. reverse-chronological) serving as the prediction target. All samples are originally in chronological order. To create balanced labels, we randomly reverse the chronology for half of the samples. Specifically, we remove the time point index for each recording, labeling them as *X* _*ϕ*1_, *X* _*ϕ*2_, *X*_ψ1_, and *X*_ψ2_. The prediction target *Y* is then defined as:

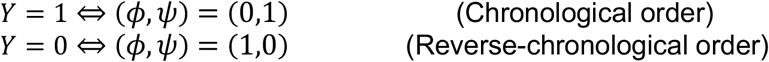

Analogous to the treatment response prediction task, we use (𝒢 _ψ_ − 𝒢 _ϕ_) as input features for chronology prediction by employing our proposed FC change-based prediction framework with classification loss. The brain pattern that predicts chronology reflects what FC changes are induced by treatment. If the classifier can generalizable across all patients in both placebo and sertraline arms while fails for healthy controls, it would essentially represents the universal placebo effect, which exists even in patients without positive treatment response.

### Contrastive Variance Quotient-Based Subtyping for Drug-Specific Responsiveness

Given the lack of significant results from the treatment arm prediction model based on linear brain dimensions (STable 2), we investigate whether the variance in brain dimensions could better distinguish FC changes between the sertraline and placebo arms. Our hypothesis is that sertraline induces FC changes consists of those from both drug-specific and placebo effects, whereas the placebo response is solely due to placebo effects. Hence, we aim to isolate the drug-specific effects by identifying brain dimensions with significantly greater variance in the sertraline arm compared to the placebo arm. This can be formularized as:

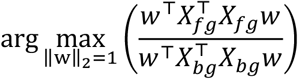

where 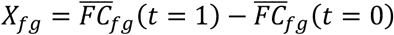 and 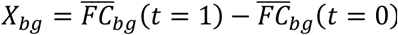 represent the differences in FC between two time points for the foreground and background data, respectively. We conceptualize the placebo effect as background noise, aiming to extract the drug-specific effect (foreground) from the overall effects in sertraline arm (foreground + background). We term this objective as the *contrastive variance quotient* (CVQ), which parallels the common spatial pattern technique used in EEG studies to separate true signals from noise^58^.

To account for natural temporal variability in FC changes, we introduce a penalty term to minimize test-retest variability, calculated as the variance of test-retest differences in FC across all subjects (pooling both treatment arms). This test-retest reliability constraint, integrated into the denominator of the CVQ, also increases the dimensionality the model can accommodate by leveraging data from both arms. We refer to this modified objective as the treatment-induced CVQ:

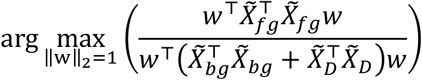

Here, 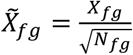 and 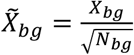 are scaled features with variances independent of sample size, where *N*_*fg*_ and *N*_*bg*_ denote the sample sizes for foreground and background data, respectively. 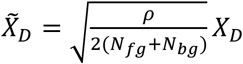 accounts for test-retest variability, with *ρ* being a hyperparameter controlling the relative importance of this penalty. The matrix *X*_*D*_ has 2(*N*_*fg*_ + *N*_*bg*_) rows corresponding to test-retest variability. With controlled feature dimensionality, the symmetric matrix 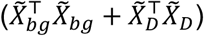 has full rank, allowing the treatment-induced CVQ to be optimized as a generalized Rayleigh quotient. In practice, we reduce feature dimensionality by selecting FC features that show potentially significant specificity to the treatment arm (uncorrected one-tailed p-value < 0.01 from the F-test).

To enhance the generalizability of findings, we integrate this CVQ-based analysis into a 5-round 20-fold cross-validation framework. In each fold, 95% of placebo-medicated patients serve as the background and 95% of sertraline-medicated patients as the foreground to identify a generalizable sertraline-specific dimension of brain change. This dimension loading is then applied to the 5% held-out placebo and sertraline arms. Aggregating results from all 20 folds provides an estimated dimension score for each subject, allowing us to assess whether the variance in dimension scores among sertraline-medicated patients is greater than that of placebo-medicated patients within the cross-validation set. To further confirm the specificity of this approach, a control analysis with placebo as the foreground and sertraline as the background is conducted. A negative result of this analysis would support the significance of the identified sertraline-specific dimension, whereas a positive result would suggest the drug-specific FC change findings may be a false positive.

Subsequently, we stratify patients in the antidepressant arm into two subtypes—those with drug-specific responsiveness and those without—based on FC change in the dimension that maximizes the treatment-induced CVQ. This subtyping stems from the observation that, while the antidepressant arm shows significantly greater variance in this dimension at the population level, not all patients have higher absolute dimension scores than those in the placebo arm (Fig. 5a). The CVQ-maximized dimension theoretically provides the highest distinguishability between the antidepressant and placebo arms, making it an ideal target of subtyping analysis. We use an F-test to assess the significance of the variance ratio between the two arms and define a threshold for the absolute dimension score in the identified dimension as:

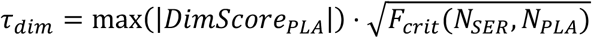

where *max*(|*DimScore*_*PLA*_|) is the highest absolute dimension score observed in the placebo arm, and *F*_*crit*_(*N*_*SER*_, *N*_*PLA*_) is the critical F-value corresponding to the sample sizes of sertraline arm and placebo arm (*F*_*crit*_ ≈ 1.5 for the EMBARC cohort at a 0.01 significance level). The critical F-value is one-tailed as our interest lies specifically in cases where patients receiving the antidepressant exhibit greater absolute dimension scores than those in the placebo arm. Since the variance is proportional to the square of the expected maximal absolute dimension score, this threshold effectively differentiates the patients with significant drug-specific responsiveness from those whose response is indistinguishable from placebo effects. Patients with absolute dimension scores above *τ*_*dim*_ are classified as having drug-specific responsiveness, while those with lower absolute scores are considered to respond similarly to placebo-treated patients.

### Generalization Analysis on Independent Cohort

Due to the absence of a placebo arm in the CAN-BIND-1 cohort and its differences from the EMBARC cohort in terms of fMRI acquisition timing and medication type, we focus on the generalization of early FC changes that are not tailored to specific prediction targets. To this end, we apply the CVQ framework to identify medication-specific FC change dimensions that are independent of clinical targets. In this analysis, the FC changes of patients within each treatment arm are treated as the foreground data, while FC changes in HCs, who received no intervention, serve as background data, allowing us to dissociate treatment-induced signal from random fluctuations and test-retest variability (Fig. 6a). The learning objective is essentially to identify FC change dimensions that maximize variance in patients while minimizing variance in HCs. This process is incorporated into a cross-validation framework to avoid overfitting. Through this approach, we aim to derive early FC change dimensions corresponding to placebo or sertraline from the EMBARC cohort, and to escitalopram from the CAN-BIND-1 cohort.

To assess generalizability, we compare each EMBARC-derived FC change dimension (placebo-induced, sertraline-induced, or sertraline-specific) to the escitalopram-induced dimension identified from the CAN-BIND-1 cohort. The sertraline-specific FC changes, derived without reference to symptom outcome, are also included. For each treatment arm, we compute the Pearson correlation between individual-level dimension scores from the CAN-BIND-1-derived escitalopram dimension and each EMBARC-derived dimension. The sertraline arm serves as an independent cohort for evaluating the correspondence between placebo- and escitalopram-derived dimensions, as it is not involved in the derivation of either. Similarly, the placebo arm serves as an independent comparison group for testing the alignment between sertraline- and escitalopram-induced dimensions. Finally, for assessing the specificity of the sertraline-specific dimension, the placebo arm is used as a negative control, where no drug-specific effects are expected.

## Data Availability

The EMBARC cohort is publicly available through the National Institute of Mental Health Data Archive (NDA) (https://nda.nih.gov/edit_collection.html?id=2199). The CAN-BIND-1 cohort is available under a data use agreement with Brain-CODE, based at the Ontario Brain Institute (https://www.braincode.ca/content/canadian-biomarker-integration-network-depression-can-bind-0).

## Code Availability

The analyses were implemented in MATLAB R2022b, and the code will be made available upon the publication of this paper.

## Acknowledgements

This work was supported by NIH grant nos. R01MH129694, R21MH130956, R21AG080425, Alzheimer’s Association Grant (AARG-22-972541), and Lehigh University FIG (FIGAWD35) and CORE grants. Portions of this research were conducted on Lehigh University’s Research Computing infrastructure partially supported by NSF Award 2019035. G.A.F. was also supported by philanthropic funding and NIH grant nos. R01MH132784 and R01MH125886, and grants from the One Mind - Baszucki Brain Research Fund, the SEAL Future Foundation, and the Brain and Behavior Research Foundation. We would like to acknowledge the individuals and organizations that have made data available for this research, including CAN-BIND, the Ontario Brain Institute, the Brain-CODE platform, and the government of Ontario.

## Financial Disclosures

G.A.F. received monetary compensation for consulting work for SynapseBio AI and owns equity in Alto Neuroscience. A.E. reports salary and equity from Alto Neuroscience and equity in Akili Interactive and Mindstrong Health. C.J.K reports equity from Alto Neuroscience. The remaining authors declare no competing interests.

## Notes

### Competing Interest Statement

G.A.F. received monetary compensation for consulting work for SynapseBio AI and owns equity in Alto Neuroscience. C.J.K reports equity from Alto Neuroscience. The remaining authors declare no competing interests. C.B.N. is a consultant for ANeuroTech (division Anima BV), Janssen Research and Development, BioXcel Therapeutics, Engrail Therapeutics, Clexio Biosciences LTD, EmbarkNeuro, Galen Mental Health LLC, GoodCap Pharmaceuticals, ITI Inc, Lucy Scientific Discovery, Relmada Therapeutics, Sage Therapeutics, Senseye Inc, Precisement Health, Autobahn Therapeutics Inc, EMA Wellness, Skyland Trail, Denovo Biopharma, and the Brain & Behavior Research Foundation. C.B.N. owns the following patents: Method and devices for transdermal delivery of lithium (US 6,375,990B1); Method of assessing antidepressant drug therapy via transport inhibition of monoamine neurotransmitters by ex vivo assay (US 7,148,027B2); and Compounds, Compositions, Methods of Synthesis, and Methods of Treatment (CRF Receptor Binding Ligand) (US 8,551, 996 B2). C.B.N. owns stock in Corcept Therapeutics, EMA Wellness, Precisement Health, Relmada Therapeutics, Signant Health, Galen Mental Health LLC, and Senseye Inc.

