## Supplementary Material for "Early Brain Functional Connectivity Changes Induced by Antidepressants and Placebo"

### Supplementary Information

**SFig. 1** Conceptual design of the chronology prediction task
**SFig. 2** Characteristics of one-week treatment-induced VPT score change in MDD patients (part 1)
**SFig. 3** Characteristics of one-week treatment-induced VPT score change in MDD patients (part 2)
**SFig. 4** Permutation testing for placebo response prediction
**SFig. 5** Correlations between placebo treatment response prediction score with VPT score change, MDD severity measures, and cognitive task performance
**SFig. 6** Correlations between placebo treatment response prediction score with childhood trauma history, personality traits, and mood and anxiety symptoms
**SFig. 7** Correlations between actual placebo treatment response MDD severity measures and cognitive task performance
**SFig. 8** Correlations between actual placebo treatment response with childhood trauma history, personality traits, and mood and anxiety symptoms
**SFig. 9** Significance and stability of the sertraline-specific FC change dimension
**SFig. 10** FC features that distinguish SER-specific and SER-nonspecific subgroups
**SFig. 11** Pre-treatment FC-based SER-specificity prediction
**SFig. 12** Comparison of distributions of SC features between SER-specific and SER-nonspecific subgroups
**SFig. 13** Distribution comparison of clinical measures between SER-specificity subgroups
**SFig. 14** Cross-validation generalizability of early FC changes induced by each medication
**SFig. 15** CONSORT flow diagram of the EMBARC clinical trial
**SFig. 16** Improved SNR in target-predictive brain dimensions

**STable 1** Sertraline Response Prediction Performance (r-values)
**STable 2** Received Treatment Prediction Performance (accuracy)

**Supplementary Methods**

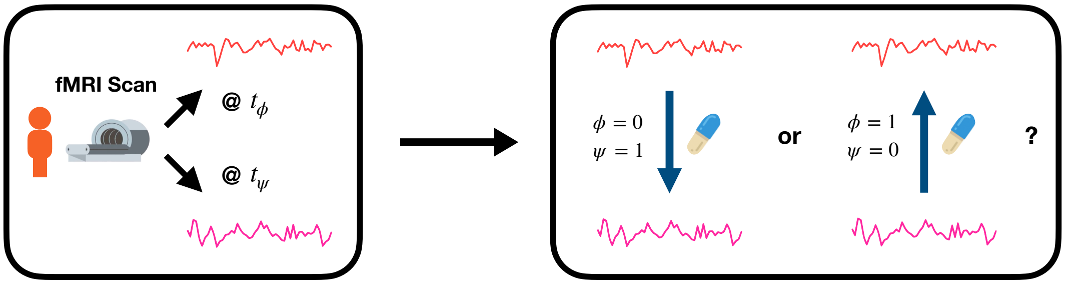


**SFig. 1** Conceptual design of the chronology prediction task. For each subject, resting-state fMRI scans are collected at two time points: pre-treatment and post-initiation. The time point indices (0 for pre-treatment, 1 for mid-treatment) are then anonymized as generic labels ($\phi$, $\psi$) to blind the model to their true order. The model is trained to predict which scan in a within-subject pair occurred earlier. To achieve high performance, the model must detect a consistent pattern of FC changes shared across all medicated patients, thereby capturing a universal placebo effect.

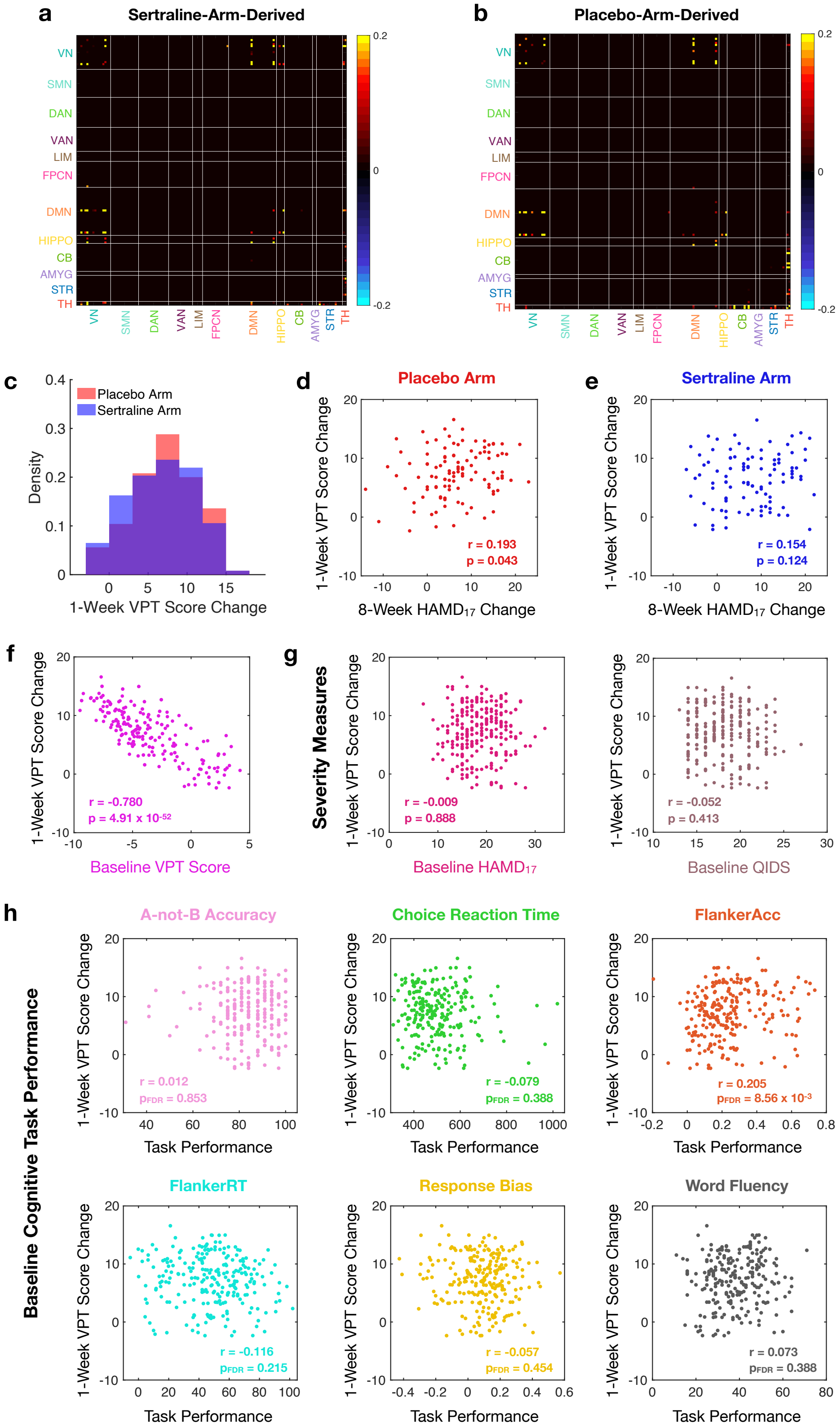
a
Sertraline-Arm-Derived
b
Placebo-Arm-Derived
VN
SMN
DAN
VAN
LIM
FPCN
DMN
HIPPO
CB
AMYG
STR
TH
0.2
0
-0.2
c
Placebo Arm
Sertraline Arm
Density
1-Week VPT Score Change
d
Placebo Arm
r = 0.193
p = 0.043
e
Sertraline Arm
r = 0.154
p = 0.124
1-Week VPT Score Change
8-Week HAMD17 Change
f
r = -0.780
p = 4.91 x 10-52
Baseline VPT Score
g
Severity Measures
r = -0.009
p = 0.888
Baseline HAMD17
r = -0.052
p = 0.413
Baseline QIDS
h
Baseline Cognitive Task Performance
A-not-B Accuracy
r = 0.012
pFDR = 0.853
Choice Reaction Time
r = -0.079
pFDR = 0.388
FlankerAcc
r = 0.205
pFDR = 8.56 x 10-3
FlankerRT
r = -0.116
pFDR = 0.215
Response Bias
r = -0.057
pFDR = 0.454
Word Fluency
r = 0.073
pFDR = 0.388
Task Performance

**SFig. 2** Characteristics of one-week treatment-induced VPT score change in MDD patients (part 1). **a-b** ROI-level FC importance for chronology prediction (universal placebo effect) derived from sertraline arm and placebo arm. Feature weights of all contributing FC features are positive, suggesting a global activation of the VPT system following one-week treatment. The predictive patterns derived from sertraline and placebo arms are highly similar (cosine similarity = 0.813), demonstrating the reliability of biomarker findings. **c** Distributions of one-week VPT score change in medicated MDD patients. No significant distribution difference is observed between treatment arms (t-test: $p = 0.446$). **d-e** Correlation between one-week VPT score change and full-course treatment response in sertraline arm and placebo arm. This correlation is statistically significant for the sertraline arm but weaker and non-significant for the placebo arm, reinforcing that VPT score change is a placebo effect. **f** Correlation between one-week VPT score change and baseline VPT score. **g** Correlations between one-week VPT score change and baseline MDD severity measures. **h** Correlations between one-week VPT score change and baseline cognitive task performance, including accuracy of the A-not-B task, choice reaction time, accuracy and reaction time of flanker test (FlankerAcc and FlankerRT), response bias of the probabilistic reward task, and valid word number in the word fluency task. p-values in **c-g** are uncorrected. p-values in **h** are FDR-corrected to validate the potentially significant correlation with FlankerAcc. Correlation coefficients in **d-g** are Pearson-based, while the correlation coefficients in **h** are Spearman-based to account for the skewed distributions of task performance metrics.

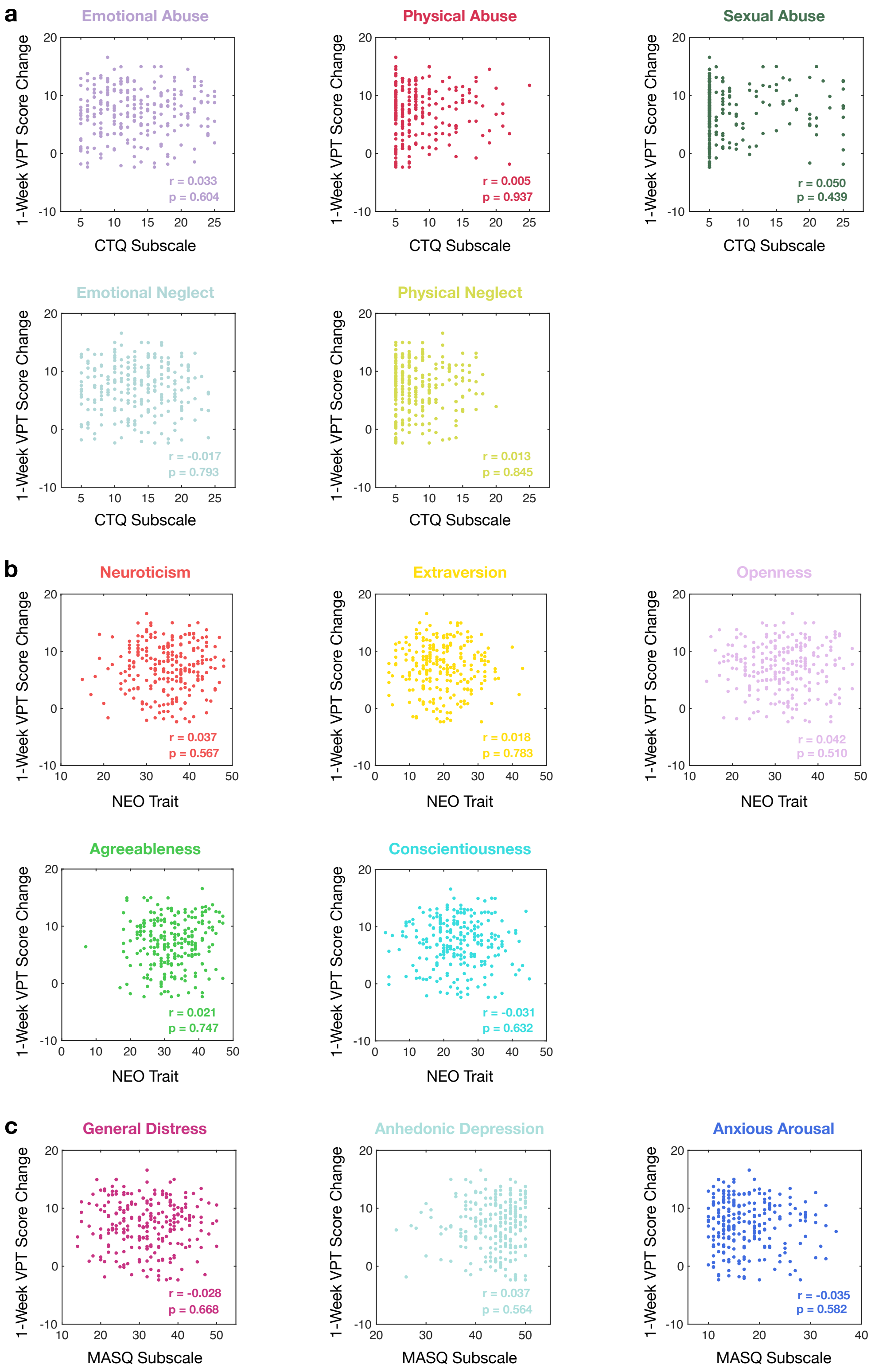


**SFig. 3** Characteristics of one-week treatment-induced VPT score change in MDD patients (part 2). **a** Correlations with Childhood Trauma Questionnaire (CTQ) subscales. **b** Correlations with NEO personality traits. **c** Correlations with Mood and Anxiety Symptom Questionnaire (MASQ) subscales. Correlation coefficients are Pearson-based. p-values are uncorrected.

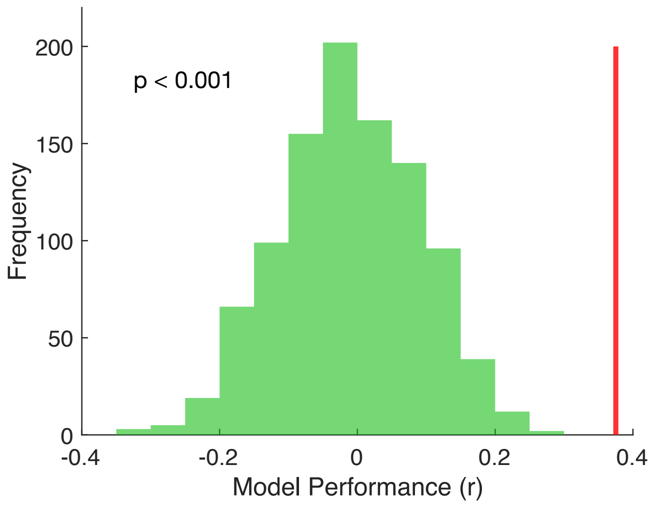


**SFig. 4** Permutation testing for placebo response prediction. The correlation coefficient (r-value) between actual and predicted placebo response is used as the model performance metric (see Fig. 3a for permutation testing of $R^2$). The red vertical line indicates the actual performance ($r = 0.375$). The green distribution shows the model performance with permuted data in 1000 trials. The permutation is conducted by assigning actual data of placebo response to random subjects.

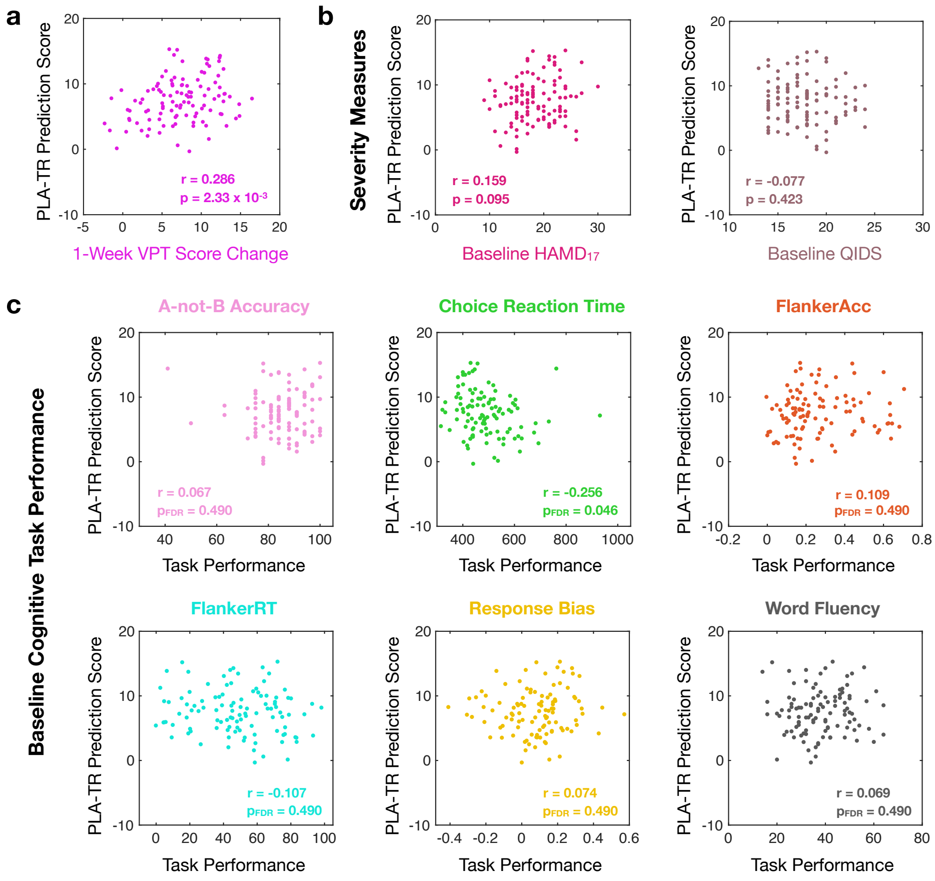


**SFig. 5** Correlations between placebo treatment response (PLA-TR) prediction score with VPT score change, MDD severity measures, and cognitive task performance. **a** With one-week VPT score change. **b** With baseline MDD severity measures. **c** With baseline cognitive task performance. p-values in **a-b** are uncorrected. p-values in **c** are FDR-corrected to validate the potentially significant correlation with choice reaction time. Correlation coefficients in a-**b** are Pearson-based, while the correlation coefficients in **c** are Spearman-based to account for the skewed distributions of task performance metrics.

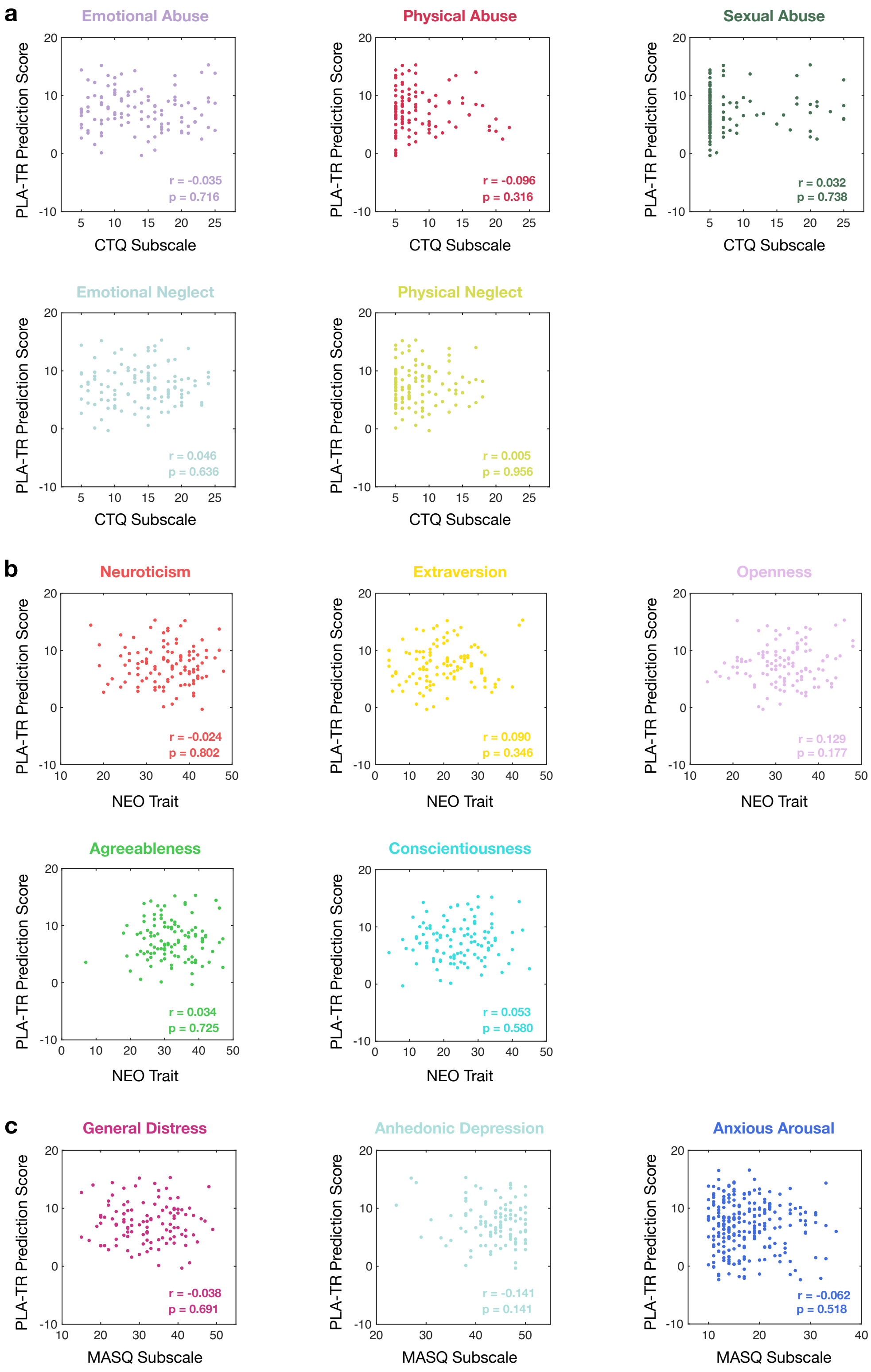


**SFig. 6** Correlations between placebo treatment response (PLA-TR) prediction score with childhood trauma history, personality traits, and mood and anxiety symptoms. **a** With Childhood Trauma Questionnaire (CTQ) subscales. **b** With NEO personality traits. **c** With Mood and Anxiety Symptom Questionnaire (MASQ) subscales. Correlation coefficients are Pearson-based. p-values are uncorrected.

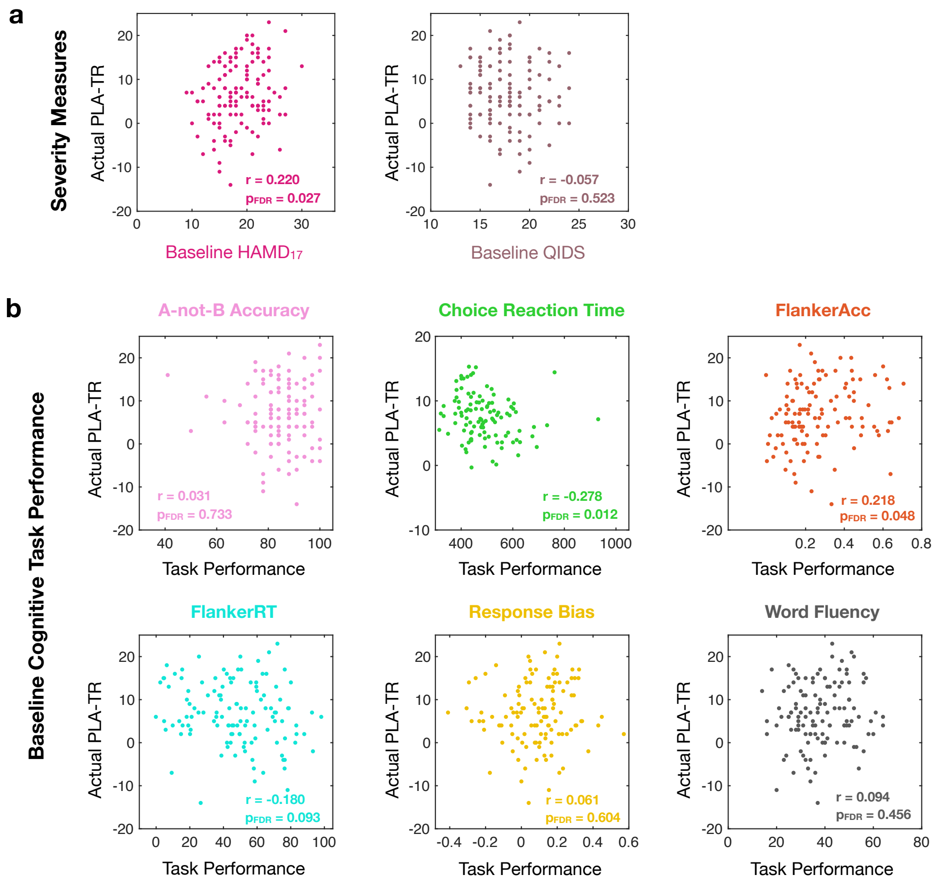


**SFig. 7** Correlations between actual placebo treatment response (PLA-TR) MDD severity measures and cognitive task performance. **a** With baseline MDD severity measures. **b** With baseline cognitive task performance. p-values are FDR-corrected to validate the potentially significant correlations. Correlation coefficients in **a** are Pearson-based, while the correlation coefficients in **b** are Spearman-based to account for the skewed distributions of task performance metrics.

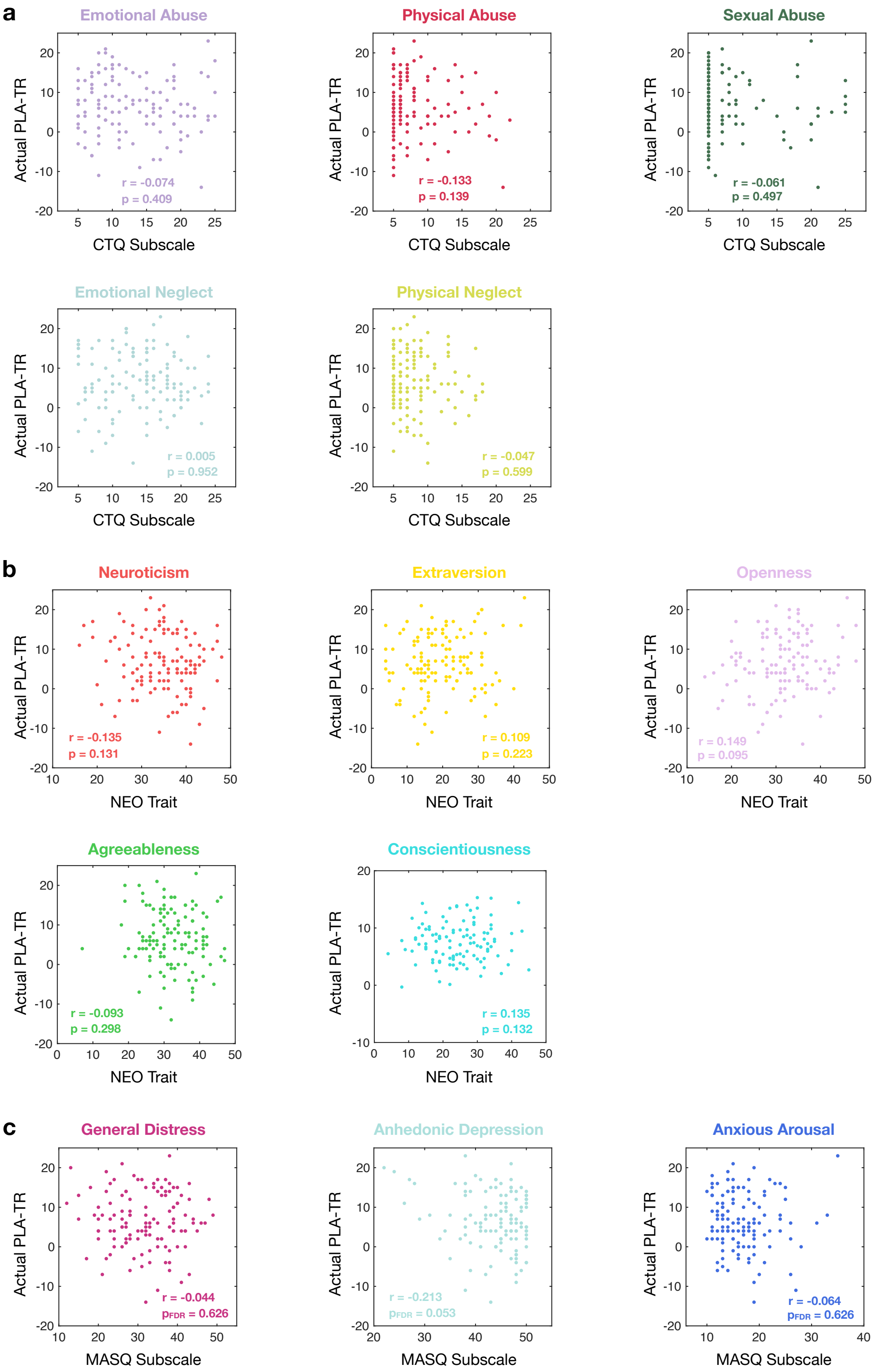


**SFig. 8** Correlations between actual placebo treatment response (PLA-TR) with childhood trauma history, personality traits, and mood and anxiety symptoms. **a** With Childhood Trauma Questionnaire (CTQ) subscales. **b** With NEO personality traits. **c** With Mood and Anxiety Symptom Questionnaire (MASQ) subscales. Correlation coefficients are Pearson-based. p-values are uncorrected.

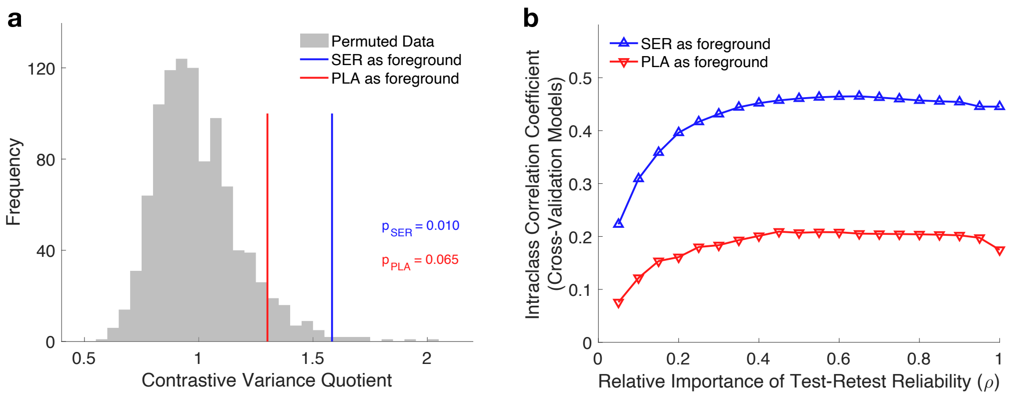


**SFig. 9** Significance and stability of the sertraline-specific FC change dimension. **a** Permutation test for the contrastive variance quotient metrics in cross-validation of the sertraline-specific and “placebo-specific” dimensions. **b** Introducing a test-retest reliability term to the background data helps the contrastive learning approach identify more stable dimensions.

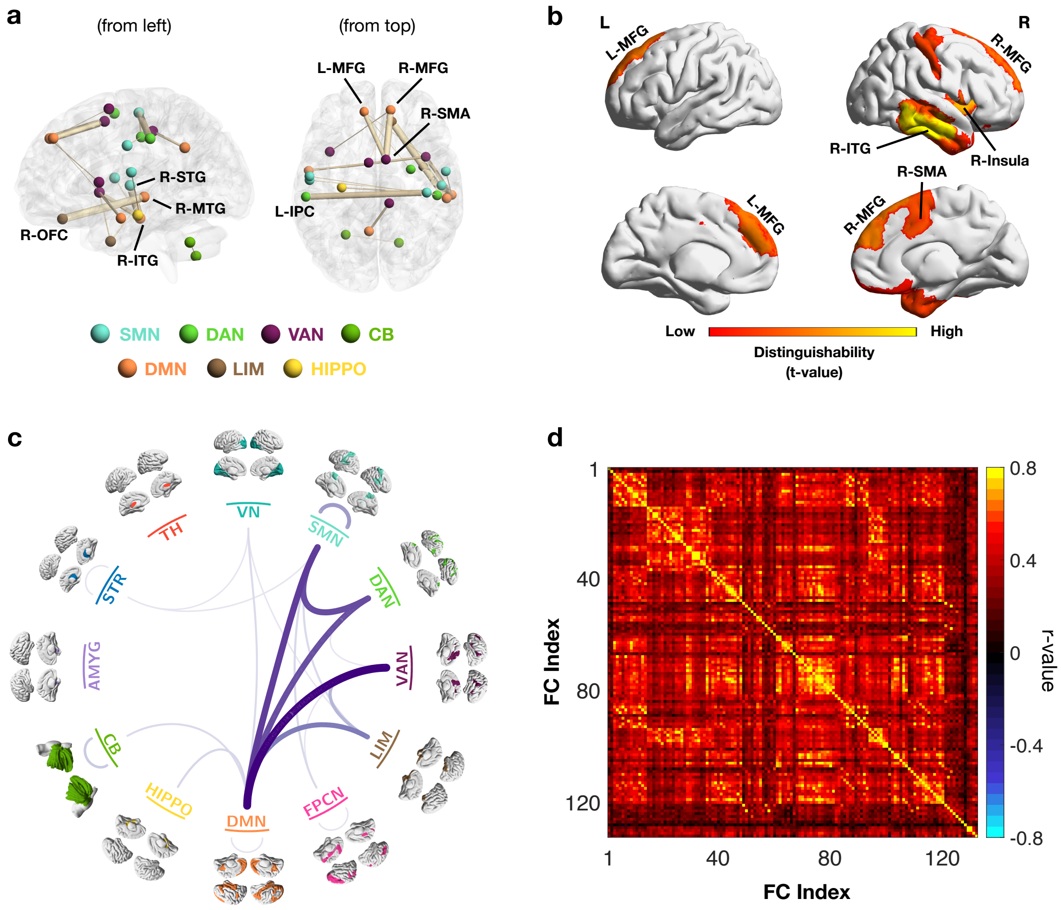


**SFig. 10** FC features that distinguish SER-specific and SER-nonspecific subgroups. **a** The FCs with the highest distinguishability across SER-specificity subgroups include the connections between the right orbital frontal cortex (R-OFC) and the right middle temporal gyrus (R-MTG), between the right superior temporal gyrus (R-STG) and the right inferior temporal gyrus (R-ITG), between the right supplementary motor area (R-SMA) and the right medial frontal gyrus (R-MFG), between the left inferior parietal cortex (L-IPC) and the right postcentral gyrus, and between the left MFG (L-MFG) and R-SMA. Thickness of connections represents effect size. **b** The brain regions with the highest distinguishability include R-ITG, the right insula, R-MFG, L-MFG, and R-SMA. The top 10 regions are displayed for clarity, which do not involve subcortical areas. **c** Important network-level connections for differentiating SER-specificity subgroups. DMN emerges as a key network, with its connections with VAN, DAN, and SMN showing the greatest difference between subgroups. The SMN-DAN connection also exhibits high distinguishability. **d** Intercorrelations across subgroup-distinguishing FC features. The global intercorrelation suggests the involving FC features form a unified dimension. Only FC features with statistically significant (FDR-corrected) difference across subgroups are included in the intercorrelation analysis.

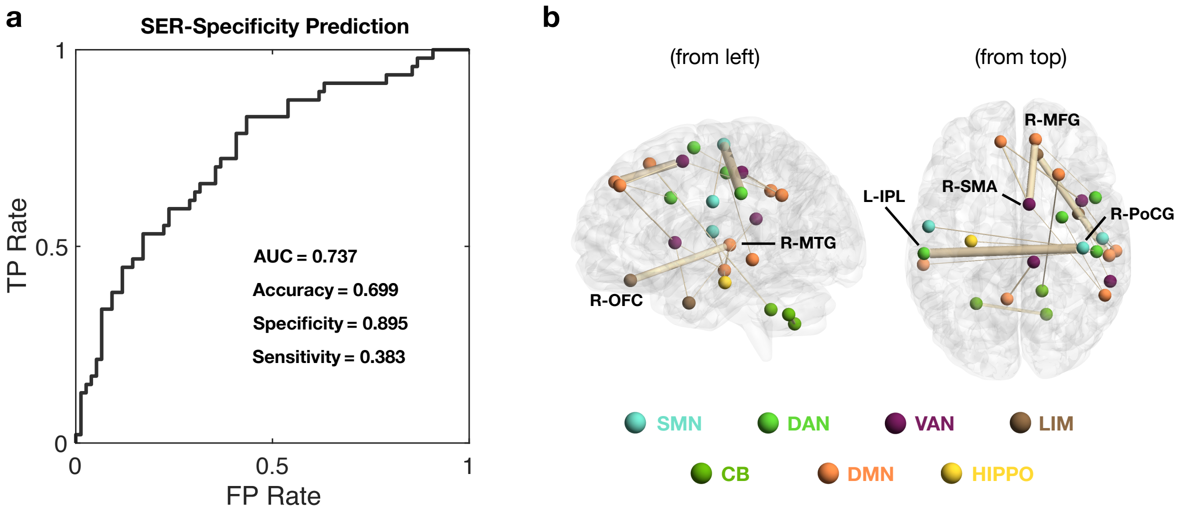


**SFig. 11** Pre-treatment FC-based SER-specificity prediction. **a** Prediction performance. The classification model is developed using logistic LASSO regression. Based on the subgrouping result (Fig. 5a), 47 patients are labeled as SER-specific (38%) and 76 as SER-nonspecific (62%). When achieving the best performance, the model made highly conservative prediction for SER-specific patients, resulting in high specificity and low sensitivity. Consequently, while the model fails to detect a majority of SER-specific patients, the predicted SER-specific patients are with high confidence level and may benefit significantly more from sertraline than placebo. This demonstrates the enhanced clinical utility of the SER-specificity subgrouping with its pre-treatment FC biomarker. **b** Brain signature. Overall, the predictive FC features are sparse, with top 3 features dominated the contribution. The most important feature is the FC between the left inferior parietal lobe (L-IPL) and the right postcentral gyrus (R-PoCG), followed by the FC between the right supplementary motor area (R-SMA) and the right superior medial frontal gyrus (R-MFG), and the one between the right superior orbitofrontal cortex (R-OFC) and the right middle temporal gyrus (R-MTG). The top 20 connections are displayed for clarity.

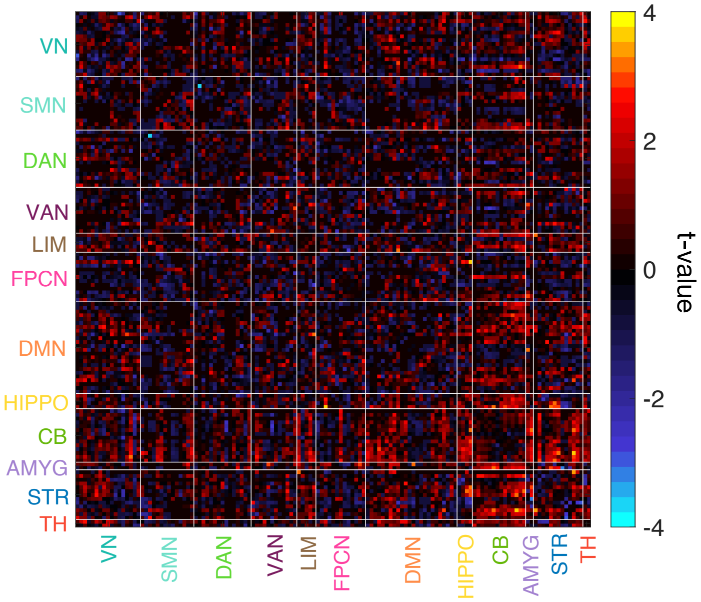


**SFig. 12** Comparison of distributions of SC features between SER-specific and SER-nonspecific subgroups. No SC features exhibit significant difference after FDR correction (*t*-test, $p_{FDR} > 0.05$).

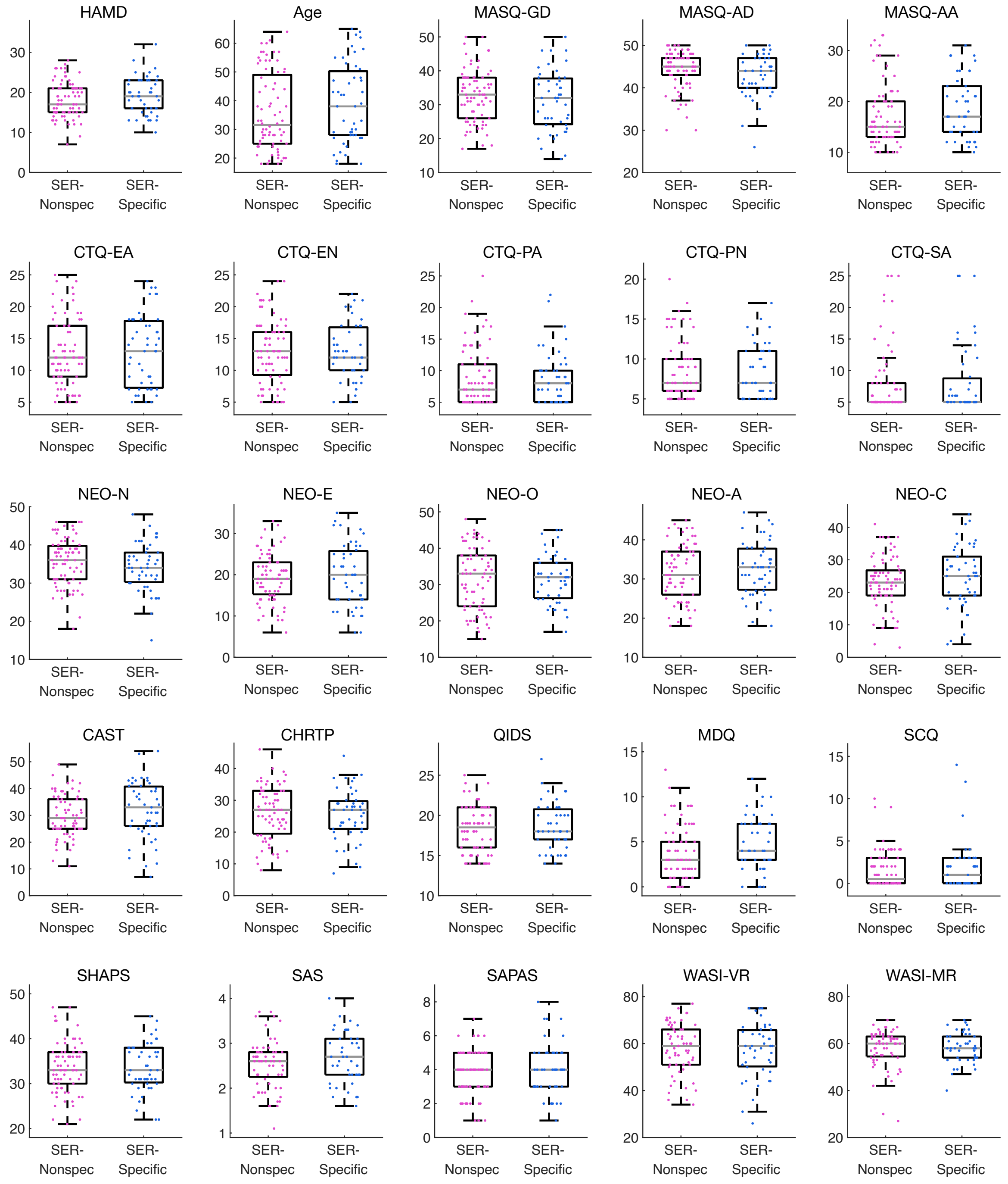


**SFig. 13** Distribution comparison of clinical measures between SER-specificity subgroups. No group difference is significant after FDR correction. HAMD: 17-item Hamilton Depression Rating Scale. MASQ (Mood and Anxiety Symptom Questionnaire): GD (general distress), AD (anhedonic depression), AA (anxious arousal). CTQ (Childhood Trauma Questionnaire): EA (emotional abuse), EN (emotional neglect), PA (physical abuse), PN (physical neglect), SA (sexual abuse). NEO personality traits: N (neuroticism), E (extraversion), O (openness), A (agreeableness), C (conscientiousness). CAST: Childhood Autism Spectrum Test. CHRTP: Concise Health Risk Tracking Propensity. QIDS: Quick Inventory of Depressive Symptomatology. MDQ: Mood Disorder Questionnaire. SCQ: Social Communication Questionnaire. SHAPS: Snaith-Hamilton Pleasure Scale. SAS: Riker Sedation-Agitation Scale. SAPAS: Standardised Assessment of Personality – Abbreviated Scale. WASI (Wechsler Abbreviated Scale of Intelligence): VR (vocabulary reasoning), MR (matrix reasoning).

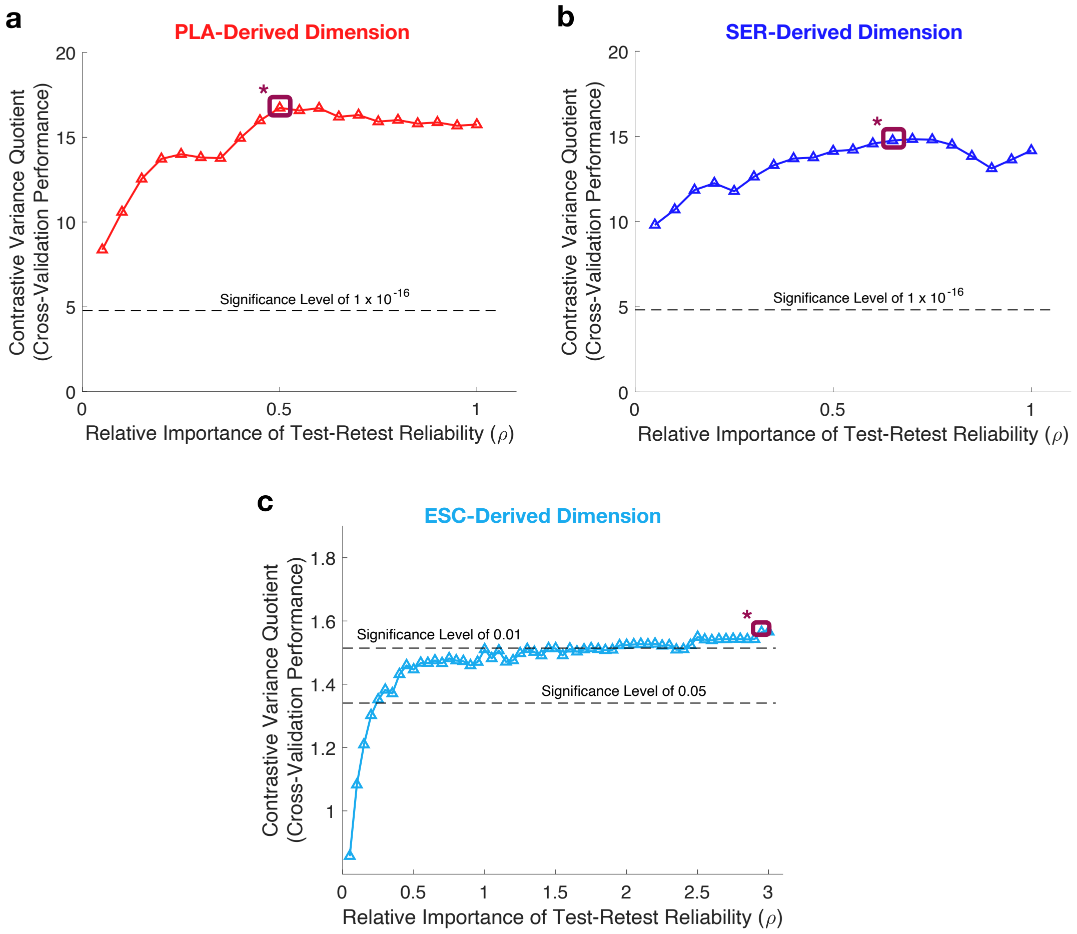


**SFig. 14** Cross-validation generalizability of early FC changes induced by each medication. **a** Placebo-arm-derived. **b** Sertraline-arm-derived. **c** Escitalopram-arm-derived (dataset 2). The identified dimension exhibits significant specificity to treatment, demonstrating the corresponding FC changes are not due to natural fluctuation and noise. Brown starred boxes indicate the operating points. The significance levels are for the one-sided hypotheses (variance: Patients > HCs).

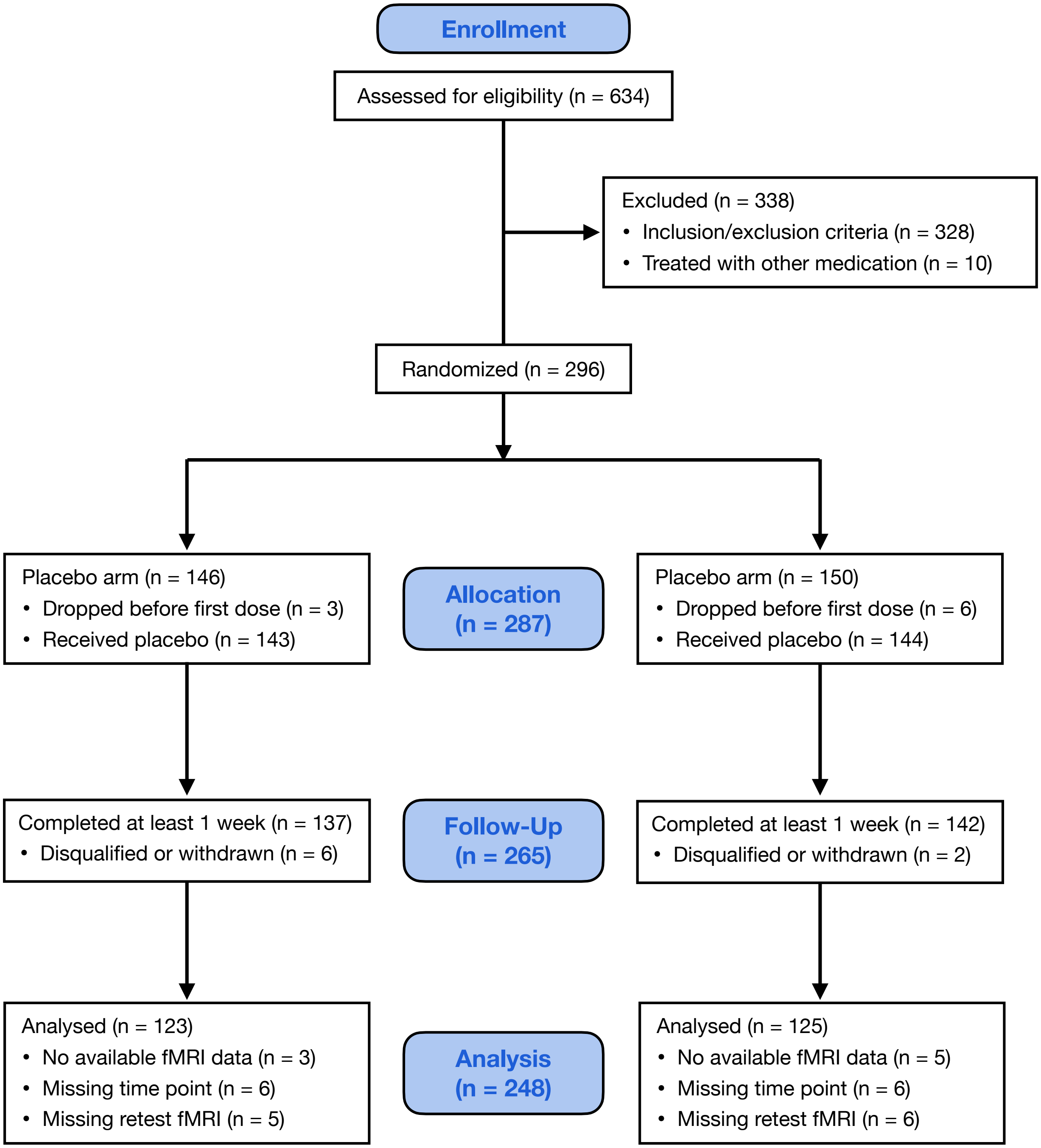


**SFig. 15** CONSORT flow diagram of the EMBARC clinical trial. The flow diagram shows the number of MDD patients who were randomized to treatment, completed at least 1 week of treatment, and had valid fMRI data available for the analyses.

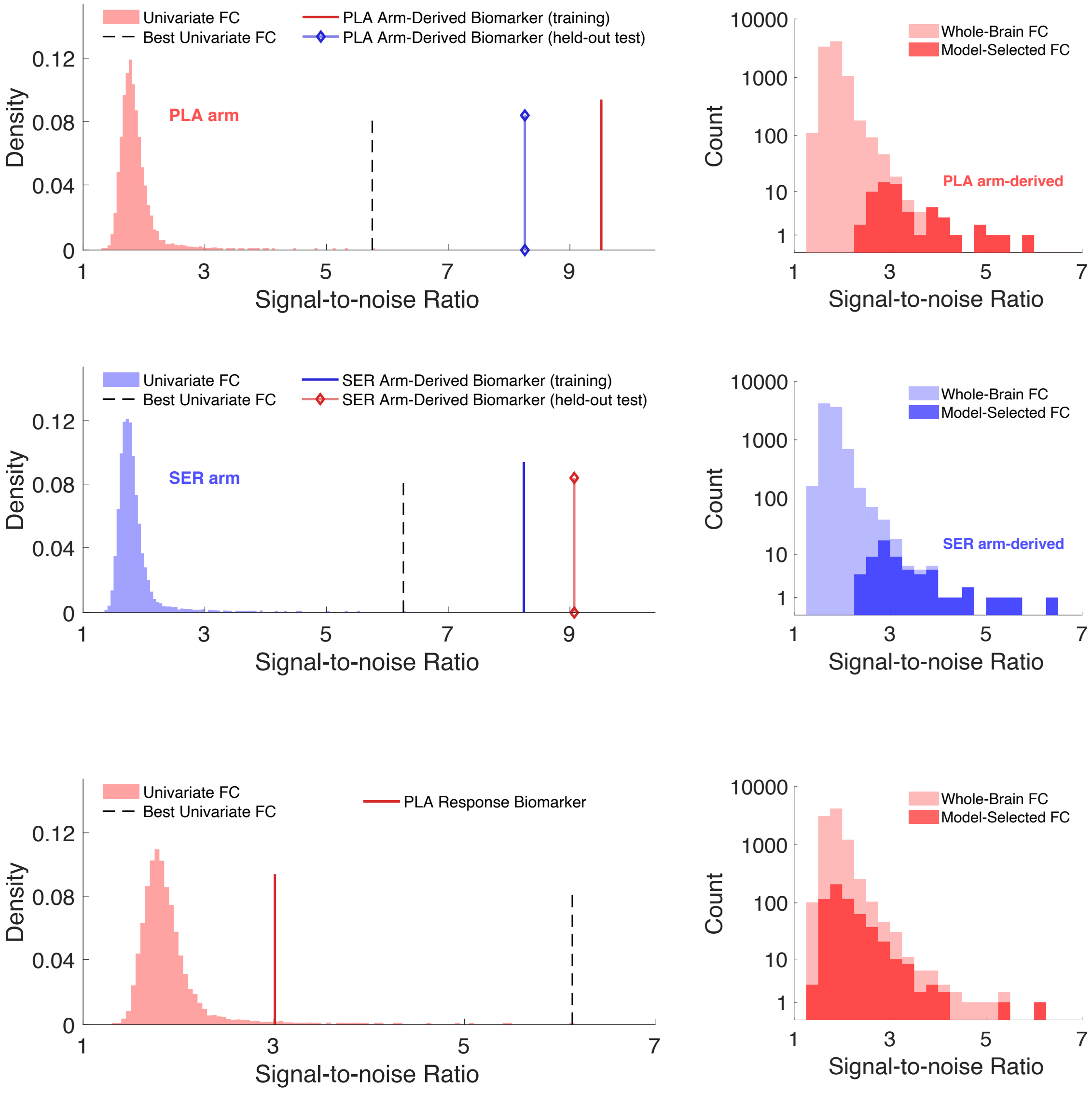


**SFig. 16** Improved SNR in target-predictive brain dimensions. Compared with univariate FC features, brain dimensions exhibit consistently higher signal-to-noise ratio (SNR) in both training and test samples. In the chronology prediction task, brain dimensions outperform even the best univariate FC feature in SNR, with the model favoring FCs that inherently have high SNR. In the response prediction task, brain dimensions show higher SNR than most univariate FCs, though lower than the top individual feature, likely due to limited target relevance among high-SNR FCs. Nevertheless, brain dimensions achieve higher SNR than the average of selected FCs while optimizing target relevance.

**STable 1** Sertraline Response Prediction Performance (r-values)

| $\lambda_{pred}$ \ $\lambda_{dim}$ | $1 \times 10^{-5}$ | $2 \times 10^{-5}$ | $5 \times 10^{-5}$ | $1 \times 10^{-4}$ | $2 \times 10^{-4}$ | $5 \times 10^{-4}$ | $1 \times 10^{-3}$ |
|---|---|---|---|---|---|---|---|
| $1 \times 10^{-5}$ | 0.102 | 0.075 | 0.010 | 0.013 | -0.008 | -0.118 | N/A |
| $2 \times 10^{-5}$ | 0.106 | 0.056 | -0.018 | 0.014 | -0.045 | -0.041 | N/A |
| $5 \times 10^{-5}$ | 0.099 | 0.056 | 0.018 | 0.025 | -0.006 | N/A | N/A |
| $1 \times 10^{-4}$ | 0.092 | -0.001 | 0.017 | -0.019 | -0.104 | N/A | N/A |
| $2 \times 10^{-4}$ | 0.035 | -0.116 | -0.072 | -0.156 | -0.134 | N/A | N/A |
| $5 \times 10^{-4}$ | -0.095 | -0.132 | -0.198 | -0.237 | N/A | N/A | N/A |
| $1 \times 10^{-3}$ | N/A | N/A | N/A | N/A | N/A | N/A | N/A |

Prediction model is developed using the SNR-constrained FC change-based machine learning framework. Prediction performance is evaluated as the Pearson correlation between actual and predicted sertraline response. $\lambda_{dim}$ and $\lambda_{dim}$ are hyperparameters controlling the sparsity of brain dimension composition and predictive feature weights (**Methods**). N/A indicates that the corresponding hyperparameter combinations resulted in all-zero models in cross-validation folds, suggesting the sparsity is overly high. For all hyperparameter combinations, $R^2 < 0$.

**STable 2** Received Treatment Prediction Performance (accuracy)

| $\lambda_{dim}$ / $\lambda_{pred}$ | 1 x $10^{-5}$ | 2 x $10^{-5}$ | 5 x $10^{-5}$ | 1 x $10^{-4}$ | 2 x $10^{-4}$ | 5 x $10^{-4}$ | 1 x $10^{-3}$ |
|---|---|---|---|---|---|---|---|
| 1 x $10^{-5}$ | 0.512 | 0.508 | 0.532 | 0.524 | 0.528 | 0.488 | 0.524 |
| 2 x $10^{-5}$ | 0.544 | 0.536 | 0.540 | 0.528 | 0.532 | 0.488 | 0.528 |
| 5 x $10^{-5}$ | 0.540 | 0.544 | 0.528 | 0.516 | 0.532 | 0.528 | 0.528 |
| 1 x $10^{-4}$ | 0.548 | 0.552 | 0.548 | 0.524 | 0.512 | 0.532 | 0.528 |
| 2 x $10^{-4}$ | 0.552 | 0.556 | 0.548 | 0.548 | 0.524 | 0.540 | 0.528 |
| 5 x $10^{-4}$ | 0.544 | 0.552 | 0.548 | 0.524 | 0.536 | 0.516 | 0.528 |
| 1 x $10^{-3}$ | 0.552 | 0.548 | 0.552 | 0.512 | 0.520 | 0.524 | 0.528 |

Prediction model is developed using the SNR-constrained FC change-based machine learning framework. The model is trained to predict whether a patient received sertraline or placebo based on one-week FC change features. Performance is evaluated as the prediction accuracy. $\lambda_{dim}$ and $\lambda_{dim}$ are hyperparameters controlling the sparsity of brain dimension composition and predictive feature weights (**Methods**).

#### Supplementary Methods

##### Diffusion MRI Acquisition

The same scanners used for fMRI acquisition were employed for diffusion MRI data collection in both cohorts. For the EMBARC cohort, diffusion MRI parameters varied by clinical site: Columbia University used a repetition time of 6 msec, echo time of 2.4 msec, 9° flip angle, and a 5-minute scan duration; Massachusetts General Hospital used 2.3 msec repetition time, 2.54 msec echo time, 9° flip angle, and 4.3-minute duration; University of Texas Southwestern Medical Center used 8 msec repetition time, 3.7 msec echo time, 12° flip angle, and 4.24-minute duration; University of Michigan used 8.1 msec repetition time, 3.7 msec echo time, 12° flip angle, and 5.29-minute duration.

In the CAN-BIND-1 study, DTI data were collected using a single-shot spin-echo echo planar imaging sequence[1]. Diffusion sensitizing gradients were applied in 31 non-collinear directions (b = 1000 s/mm²) and 6 volumes with b = 0 s/mm². Despite some protocol variations[1] across sites, key parameters were consistent, including: 94 msec echo time, 90° flip angle, 64 × 64 matrix size, and voxel dimensions of 2.5 mm × 2.5 mm × 2.5 mm.

##### Diffusion MRI Preprocessing

Diffusion MRI images with b-values less than 100 s/mm² were designated as b=0 images. Denoising was then performed using MRtrix3's method[2] with a 5-voxel window. Subsequently, B1 field inhomogeneity was corrected using the dwibiascorrect function from MRtrix3 with the N4 algorithm[3]. The mean intensity of DWI series was standardized to match the b=0 image intensity across each scan. FSL's eddy tool corrected for head motion and Eddy correction[4], utilizing a q-space smoothing factor of 10, 5 iterations, and 1000 voxel-based hyperparameter estimation. First- and second-level linear models were used to adjust for Eddy-induced spatial distortions, with q-space coordinates assigned to their respective shells. Field offset was separated from subject movement, followed by outlier replacement using Eddy's outlier replacement method[5]. Data were organized by slices, retaining those with at least 250 intracerebral voxels, and outliers deviating by more than 4 standard deviations were replaced with imputed values.

##### Structural Connectivity Calculation

Structural connectivity (SC) was computed using DSI studio software (https://dsi-studio.labsolver.org). First, a white matter mask was generated with thresholding. The spin distribution function for each voxel within the masked image was then estimated through generalized q-sampling imaging (GQI) reconstruction[60] in T1-weighted space, with a diffusion sampling length ratio of 1.25. Tensor metrics were derived from DWI data with a b-value lower than 1750 s/mm². Whole brain fibers were reconstructed using a modified streamline deterministic tracking algorithm[61], with augmented tracking strategies[62] and whole brain seeding. Key parameters were set as follows: angular threshold of 45°, step size of 1 mm, fiber length extent 20–300 mm, and maximum subvoxel search seeds 1,000,000. In each iteration, the algorithm randomly selected a voxel in the brain, initiating fiber tracking in both directions until the fiber reached the brain boundary. Tracks shorter than 30 or longer than 200 mm were excluded. Finally, SC was derived by counting fibers connecting each pair of regions using the same brain atlas as in the FC calculation.
